# EvoSeq-ML: Advancing Data-Centric Machine Learning with Evolutionary-Informed Protein Sequence Representation and Generation

**DOI:** 10.1101/2024.10.02.616302

**Authors:** Mehrsa Mardikoraem, Nathaniel Pascual, Joelle N. Eaves, Shramana Chatterjee, Anton Mahama, Patrick Finneran, Robert P. Hausinger, Daniel R. Woldring

## Abstract

From protein structure prediction to novel protein generation, challenging protein engineering tasks have been made possible by advancements in machine learning (ML). While largely driven by ML architecture refinements, these advancements in ML-based protein engineering campaigns have left the impact of data curation underexplored. In light of the growing wealth of labeled sequence data, data-centric advances (e.g. prioritizing improvements in ML protein engineering tools through the curation of high-quality, domain-specific training data) are increasingly preferred over model-centric advancements. Implementing datasets that accurately reflect biological complexity and diversity has been shown to improve the efficiency of training protein engineering ML tools. Here, we evaluate an ancestral sequence reconstruction (ASR)-informed data augmentation strategy for training generative and representation-learning models in protein engineering. Using ethylene-forming enzyme (EFE) as a model system, we show that variational autoencoder models trained on ancestral and near-ancestral sequence datasets generate variants with improved predicted and experimentally measured thermostability relative to variants generated from modern-sequence training data. All experimentally tested ancestral and ML-generated EFEs produced detectable ethylene, although ML-generated variants showed reduced activity relative to wild-type EFE, indicating that the approach more strongly captured stability-associated features than catalytic optimization. We further evaluated ASR-enriched sequence sets for fine-tuning ESM2 representations in endolysin and lysozyme C stability-classification tasks, where ancestral representations were competitive with modern-sequence fine-tuning in selected settings. Overall, this work supports ASR-informed data augmentation as a promising strategy for stability-oriented protein sequence generation and motivates future work to couple ancestral sequence diversity with explicit functional selection.

## Introduction

Recent advancements in machine learning (ML) highlight the critical role of data quality and diversity in model performance, shifting focus towards data-centric approaches that complement algorithmic innovations^1,2^. This evolving perspective underscores the importance of curating diverse datasets in developing effective ML models. Unlike model-centric methods that are widely accessible through open-source platforms, data-centric methods offer tailored insights to specific applications which enhances model learning capabilities beyond scoring metrics^3,4^. This trend is evident in the development of the GPT model, which evolved from focusing on architectural improvements to prioritizing data quality and data collection strategies^5^. As a result, focusing on data-centric approaches promises a profound contribution toward ML-driven problem-solving in distinct domains. These approaches have already exhibited potential in fields ranging from finance to healthcare, improving model reliability, scalability, trustworthiness, and generalizability^2,6–9^. This shift towards data-centric approaches is especially significant in areas demanding intricate analysis, like protein engineering, where such strategies could enhance the understanding of the complex protein fitness landscape^10,11^.

In protein engineering, the adoption of data-centric approaches is paramount due to the field’s unique challenges. The protein fitness landscape is complex and rugged, with minor alterations to protein sequences significantly impacting functionality and stability^12–14^. Despite advances in applying ML that have facilitated the discovery of high-fitness proteins^15,16^, these models struggle with imbalanced datasets and the scarcity of data across protein families. This concern underscores the necessity of reevaluating our strategies towards curating training data^10,11,17–20^. A focus on enhancing the diversity and quality of datasets is crucial, as it ensures that ML models are trained on data that accurately reflect the intricate biological properties of proteins. Establishing a method to provide more effective training datasets for ML models will pave the way for strategically advancing protein engineering campaigns.

Built on the foundation of evolutionary biology and molecular phylogenetics, ancestral sequence reconstruction (ASR) constructs phylogenetic trees from an alignment of extant sequences and subsequently implements substitution models to infer ancestral sequences - sequences that are typically diverse, and stable^21–24^. Therefore, we hypothesized that enriching multiple sequence alignments - typically consisting of modern sequences - with high-quality ancestral sequences would be a powerful data-centric approach to protein engineering.

Furthermore, we explored methods to exploit the inherent statistical uncertainties of ASR to dramatically expand the number of ancestral sequences from which ML models can be trained. Guided by various substitution models for mutations, ASR assigns posterior probabilities to amino acids at various positions to infer ancestral sequences at a given node in a tree. Typical ASR campaigns reconstruct ancestors by picking amino acids with the highest probability. However, sampling from a broad distribution of amino acids can generate ancestral proteins that are functional and, in some cases, more efficient or offer novel functionalities compared to the most probable ancestors^25,26^. Even when incorporating alternative amino acids with a posterior probability as low as 30%, reconstructed proteins maintained their overall function^25^. Given its robustness to generate diverse and functional libraries despite statistical uncertainties, ASR can generate up to a billion unique sequences for a protein with 15 positions that each have four high-likelihood amino acids.

Therefore, with the potential to generate largescale high-quality sequences, ASR represents a potentially impactful data-centric protein engineering approach. Prior ASR studies have shown that alternative ancestral states arising from statistical uncertainty can often retain function, motivating the use of ancestral ensembles rather than only single maximum-probability reconstructions***. Recent work by Colin Jackson’s lab has extended this concept into ML workflows by using multiple reconstructed ancestral sequences to improve protein representation learning ^27,28^. It accomplishes this by using ASR to obtain protein representations that are family specific. Their approach employs a custom transformer model (LASE) trained on ancestral sequences reconstructed by sampling the maximum *a posteriori* (MAP) characters for a given site across several hundred independent ASR instances. While this method has shown significant improvements in protein representation learning and downstream task performance, particularly demonstrating a smoother fitness landscape and computational efficiency for the phosphodiesterase family, it has limitations. In particular, sitewise MAP or marginal posterior sampling approaches do not explicitly sample from the full joint distribution of ancestral sequences. Thus, while these approaches incorporate uncertainty at individual sites, they should be distinguished from methods that directly model site-site dependencies or epistatic constraints.

Building on this foundation, we evaluate ASR-derived and near-ancestral sequence ensembles as data augmentation sets for two ML protein-engineering tasks: (1) the generation of novel protein sequences through generative modeling and (2) the enhancement of protein classification via fine-tuned language models (Figure 1). For fine-tuning, we utilized evolutionary scale modeling (ESM) protein representations^29^, comparing representations trained on two protein families, endolysin and lysozyme C, against representations fine-tuned with ancestral sequences of these two protein families. For the generative ML tasks, we implemented a variational autoencoder (VAE) using reconstructed sequence data obtained through both Bayesian (via BAliPhy) and maximum likelihood (via IQ-TREE) methods in ASR for the ethylene-forming enzyme (EFE)^30^ isolated from *Pseudomonas savastanoi* pv. *phaseolicola* PK2^31^. The structures and thermal stabilities of the ML-generated sequences were predicted using AlphaFold2^32^ and FoldX^33^, respectively. We compared sequences generated from ancestral data to those generated from modern sequences using metrics of structural stability, sequence variability, and semantic diversity. Furthermore, we provided wet-lab validation of the activity and stability of eight ancestral sequences and four EFEs generated from ancestral sequences using ML. To assess whether diversification preserved functional chemistry, we used AlphaFold3 (AF3)^34^ joint structure prediction and molecular docking to show that ancestral and ML-generated EFEs retain conserved structural and binding interactions features. Altogether, these innovations advance the use of evolutionary information in protein engineering, offering new tools for sequence generation and representation learning, and enhancing exploration of the protein fitness landscape.

**Figure 1.**
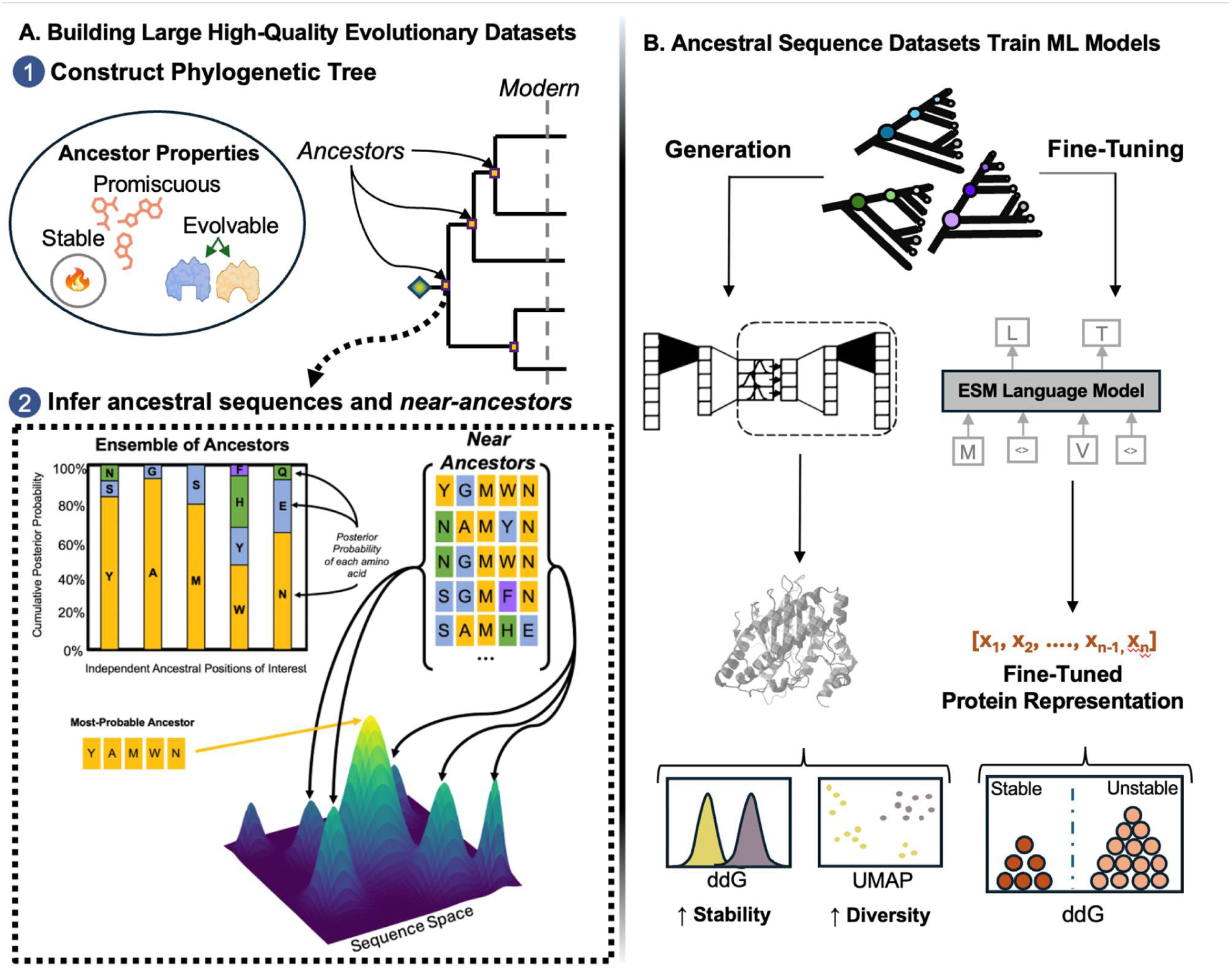
Large Ancestral Datasets are Compiled from the Evolutionary Landscape to Train ML Models A. Generation of high-quality evolutionary data. The phylogenetic tree for the family of interest and ancestral sequences are inferred from modern sequences. Ancestral sequences are often known to be stable, promiscuous (i.e. substrate specificity is relaxed), and evolvable. To generate ancestral sequences, IQ-TREE employs marginal reconstruction, which calculates the most likely state (amino acid), shown as yellow nodes, for each site (residue position number) independently. Ensembles of ancestral sequences can then be produced by sampling from other probable amino acids at individual positions of the ancestral sequences. **B. Ancestral sequences can be implemented in generative ML tasks.** The obtained evolutionary information is used as training data for sequence generation and family-specific protein sequence representations. For the former, a general VAE is trained on selected ancestral ensembles and sequences are generated by sampling from the latent space. For the latter, ESM representations were fine-tuned using ancestral sequences and evaluated based on thermostability prediction.

**Figure 2.**
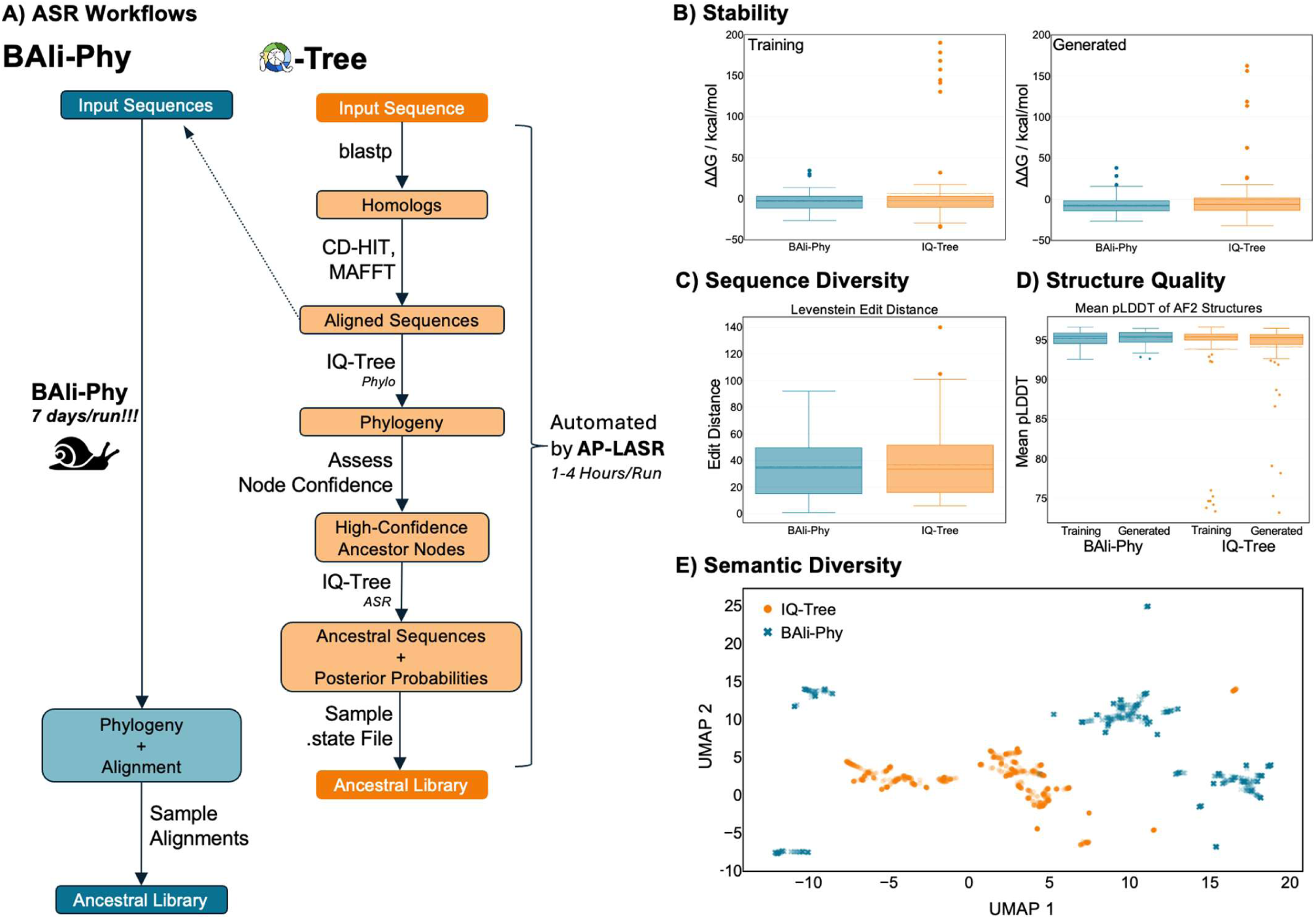
Maximum Likelihood (IQ-Tree) and Bayesian (BAli-Phy) approaches for ancestral sequence reconstruction of the EFE family are comparable in terms of thermostability, sequence variability, and structural quality. A. Visual outline of Maximum Likelihood and Bayesian workflows to curate ancestral sequences used for VAE-based generation tasks. Of note, the same alignment of extant sequences is used for both Bali-Phy and IQ-Tree workflows. B. Stability analysis using FoldX predictions of thermodynamic stability (ΔΔG, kcal/mol) between generated and training sequences suggest no statistically significant differences were observed (p-value=0.17). C. Sequence variability illustrated by minimum edit distances between training and generated sequences reveals similar diversity for both methods. D. Structural quality analysis using pLDDT scores demonstrated no significant differences between methods (p-value=0.37 for means and p-value=0.33 for IDRs). E. Similarly, semantic diversity visualized via UMAP projections of sampled generated sequences indicate comparable patterns.

## Methods

### Generative Model for Novel Protein Sequence Generation

We aimed to test whether incorporating evolutionary data into the training of generative models yields protein sequences that are both novel and structurally stable. To this end, we focused on the strain PK2 EFE and employed a VAE as our generative ML model. Subsequent computational analyses (i.e., AlphaFold^32^, FoldX^33^, UMAP visualization^35^) were conducted on the sequences generated by this model to assess their quality.

#### Ancestral Sequence Reconstruction

We extracted the evolutionary information of EFE using our AP-LASR^36^ software, a tool designed to reconstruct ensembles of ancestral proteins by leveraging the phylogenetic tree of a query protein sequence^37^. This tool has facilitated ASR by fully automating the reconstruction process from initial BLAST search, multiple sequence alignment by MAFFT^38^ to tree phylogeny predictions via IQ-TREE2^39^.

For this study, we focused on two AP-LASR outputs: maximum-probability ancestral sequences (ASR-Max), and diversity-enhanced near-ancestral sequence ensembles (ASR-Dist). ASR-Dist was generated from the sitewise marginal posterior probabilities reported in the IQ-TREE ASR.state file. For each ancestral node, amino acids with posterior probability above a defined threshold were retained as allowable states at that position, and sequences were then sampled from these allowable states to generate near-ancestral libraries. Therefore, ASR-Dist should be interpreted as a diversity-enhanced sampling of plausible near-ancestral sequence space based on marginal posterior uncertainty, rather than as exact draws from the full joint posterior distribution of ancestral sequences. Unless otherwise noted, we used a posterior-probability threshold of 0.2 based on prior evidence that ancestral proteins can be robust to moderate posterior uncertainty and because this threshold provided a practical compromise between library diversity and retention of high-probability ancestral states ^25,40^.

To evaluate how the choice of probabilistic sequence reconstruction methods influences generative model performance, we compared IQ-TREE2’s maximum likelihood approach against BAli-Phy’s Bayesian inference method. BAli-Phy uses a Bayesian inference method to simultaneously estimate sequence alignment, tree phylogeny, and model parameters, providing full posterior distributions of ancestral sequences^41^. Because BAli-Phy jointly estimates alignment, tree topology, model parameters, and ancestral states, it can account for additional sources of uncertainty relative to workflows that condition on a fixed alignment and tree. This increased uncertainty modeling comes at substantially greater computational cost.

In brief, the same multiple sequence alignment generated by AP-LASR for IQ-TREE was used to initialize eight independent Markov chain Monte Carlo (MCMC) runs in BAli-Phy. The optimal amino acid substitution model for the runs, lg08^42^, was determined using IQ-TREE’s ModelFinder^43^. After running simulations for seven days, three runs were selected for convergence by minimizing the average standard deviation of split frequencies (ASDSF) and the maximum of the standard deviation of split frequencies (MSDSF). For convergence analysis, the first 2,000 iterations of each run were discarded to account for the bias of the initial random starting guess (i.e., burn-in). The ensemble of ancestors at each ancestral node were subsequently retrieved from the BAli-Phy runs, using the MAP tree topology to define ancestor nodes. ETE4 Toolkit^44^ was used to define all ancestor nodes on the MAP tree based on each node’s set of children sequences, and Dendropy^45^ was used to sample ancestral sequences at each ancestral node for all BAli-Phy output trees. For more targeted protein generation, we sampled the sequences from four nodes representing high-stability ancestors obtained from various evolutionary timescales (Node10, Node13, Node253, and Node384, as shown in Figure S1)^37^.

#### Automated BAli-Phy replicate workflow and ASR generation

To improve robustness of alignment, topology, and ancestral sequence estimates, we implemented an automated BAli-Phy–based workflow that runs multiple independent MCMC replicates and summarizes a consistent subset for downstream ASR. The pipeline (executed on an HPC cluster via SLURM with BAli-Phy packaged in a Singularity container) launches five BAli-Phy replicates from the same input FASTA, extracts posterior values from each run after burn-in, and selects the three runs with the most similar post–burn-in median posterior values to exclude obvious outlier chains prior to downstream summarization. These selected replicates are then passed to bp-analyze to generate a consensus summary and the maximum a posteriori (MAP) tree (MAP.tree). A consensus alignment is reordered to match the MAP tree tip order (alignment-cat) and summarize-ancestors is used to generate an ASR.state file and associated outputs for downstream AP-LASR analyses. Detailed, step-by-step usage instructions (inputs/outputs, commands, and troubleshooting) are provided in the Supplementary Instructions and in the accompanying repository. All BAli-Phy tools are executed within the container, and intermediate outputs (posterior summaries, consensus alignments, and trees) are retained to facilitate troubleshooting and rerunning summary steps without repeating the full MCMC.

#### Dataset Curation

From the data generated by AP-LASR, we sampled the sequences from four nodes that represented high-stability ancestors obtained from various evolutionary timescales (Figure S1). To understand how sequence diversity in our training set would impact the robustness and generalizability of our model, two distinct datasets of ancestral protein sequences were crafted for training our ML model. The “Homogeneous Dataset” is comprised of sequences equally sampled from ancestral nodes, whereas the “Diverse Dataset” was created by passing the initial dataset through CD-Hit with a 0.9 similarity threshold^46^. The sampled sequences for each node were initially aligned using MAFFT^38^. Subsequently, the sequences from all ancestral nodes were pooled and re-aligned with MAFFT. A representative subset of sequences (1,000 sequences per data type) was randomly sampled from each dataset for both fine-tuning and generation tasks.

#### Generative Model Architecture and Training

VAE was selected for its proficiency in generating new data points that are coherent with the training data^47,48^. We employed one-hot encoding to transform the sequences into a format suitable for computational processing. Feature extraction from these one-hot encoded sequences was performed using a 1D Convolutional Neural Network (CNN) layer, allowing us to capture the local sequence patterns effectively. The architecture of our VAE was designed with a latent space dimensionality of 100, ensuring sufficient complexity to capture the nuances of protein sequence variability. Additionally, we incorporated batch normalization within the network to facilitate smoother and more stable learning dynamics. This combination of 1D CNN for feature learning and batch normalization for optimization contributed to refining the model’s ability to generate meaningful protein sequences. Novel sequences were generated by sampling from the learned latent representations after loss minimization in the validation set.

#### Evaluation

To gauge the ability of our models to generate stable and viable sequences, we utilized AlphaFold2 to verify that the predicted 3D structures of the generated EFE ancestors exhibited folding patterns similar to the wild-type structure. Randomly selected sequences analyzed by AlphaFold2 were superimposed onto wild-type structures (PDB ID: 5V2Y) using PyMol to compare folding patterns (as indicated by the RMSD between the wild-type and training sequences). Stability calculations (ddG) of these predicted structures were carried out using FoldX. This evaluation phase was crucial, as it allowed us to ascertain not just the novelty of the generated sequences but also their practical applicability in terms of structural integrity and thermal stability. We used the Dunn statistical test to compare the distribution of predicted structure stability measurements. The sequences generated by a model trained on evolutionary-reconstructed sequences were compared to those generated by a model trained on modern sequences. The comparison focused on three aspects: structural quality, sequence diversity, and semantic diversity.

To assess structural quality, we used the predicted Local Distance Difference Test (pLDDT) scores which were extracted using the alphapickle library^49^. Ranging from 0 to 100, pLDDT is used to assess the reliability of the predicted atomic positions in the protein structure. We used the Levenshtein edit distance^50,51^ (i.e., the minimum number of single-character edits required to change one string into another) to quantify the divergence between training sequences and generated sequences across different datasets. This approach allowed us to assess the sequence variability introduced by the generation process. To characterize semantic diversity, which considers the biological or chemical properties of the sequences rather than just their raw sequence differences, the sequence representations obtained via ESM language model were visualized with UMAP, a dimensional reduction technique that represents the data manifold in lower dimensions^35^.

To further probe the wet-lab results described below, 50 randomly selected “diverse” generated sequences, 50 randomly selected “homogeneous” generated sequences, and 50 randomly selected ancestral sequences were structurally modeled and docked to their substrates using AlphaFold3 to investigate potential differences in protein-ligand interactions with the wild-type EFE bound to Mn(II), 2-oxoglutarate (2OG), and L-arginine (L-Arg). The mmCIF outputs of AlphaFold3 were converted to PDB file format using BeEM^52^, and these PDB files were subsequently analyzed with the protein-ligand interaction profiler (PLIP)^53^ to index potential protein-ligand interactions. The mutation frequency at each position of the EFE and the sequence identity matrix was determined on an alignment of sequences generated by MAFFT^38^. ESPript 3.0 was used to visualize these alignments and highlight key conserved protein-ligand interactions.

### Evolutionary-Derived Protein Sequence Representation

In this section, we detail our approach to creating family-specific protein representations using fine-tuned methods. We divided the fine-tuning into two categories: regular fine-tuning, which adjusts model parameters using the InterPro protein family dataset, and evolutionary fine-tuning, which incorporates specific evolutionary data relevant to protein families to guide the tuning process. The performance of these fine-tuned representations was evaluated by comparing them against the baseline ESM2 (Evolutionary Scale Modeling)^29^ representation in a protein stability prediction task.

#### Data Processing

We fine-tuned the ESM2 for endolysin and lysozyme C, for which we obtained labeled datasets for stability prediction in FireProtDB^54^. Our data processing involved assembling three distinct unlabeled datasets to fine-tune the ESM model for each protein family: (i) a collection of InterPro-derived sequences that encompass an expanded set of modern proteins based on family affiliations^55^, (ii) a collection of the single most-probable ancestors (ASR-Max) taken from every internal node of the phylogenetic tree built with AP-LASR, and (iii) a collection of ancestral ensembles sampled from internal nodes (ASR-Dist). The ASR-Dist approach enriched the dataset with a broader spectrum of evolutionary possibilities, offering a more robust dataset that surpassed ASR-Max in both diversity and volume. This comprehensive dataset integrated extensive evolutionary insights, significantly enhancing the model’s training base. We sampled 1,000 sequences inferred from each high-quality ancestral node (which we defined as having SH-aLRT *>* 80% and ultrafast bootstrapping *>* 95%) reconstructed in ASR and removed repeat sequences. Table 1 represents dataset information for both protein families tested.

**Table 1.** Datasets Used for Fine-Tuning Task.

| Protein | Interpro ID | Interpro Seqs | ASR-MAX Seqs | ASR-Dist Seqs | FireProtDB Seqs |
| --- | --- | --- | --- | --- | --- |
| Endolysin | IPR034690 | 12K | 302 | 40K | 647 |
| Lysozyme C | IPR036328 | 9K | 452 | 36K | 338 |

#### Classification Model Training

For model fine-tuning, we employed the ESM2 model (esm2 t12 35M UR50D) trained with 35M parameters which generated 480 embedding dimensions and contained 12-layer representations. We unfroze its last two layers to adapt its learning to our specific datasets. A batch size of 32 sequences was utilized to optimize the training process, alongside the implementation of early stopping to mitigate the risk of over-fitting. Post-tuning, we extracted the embedding from each of the four datasets to further refine our approach to protein family classification, employing KNN^56^, Random Forest^57^, and XGBoost^58^ algorithms to assess the model’s predictive performance within each representation derived from distinct fine-tuning methods.

#### Evaluation of Classification Models

The prediction task was stability classification (i.e., determining if a given sequence was stable with a ΔΔ*G <* −0.5 kcal/mol or unstable with a ΔΔ*G >* 0.5 kcal/mol). Precision, recall, balanced accuracy, Area Under the Curve (AUC), and the F1 score were calculated for each protein family (endolysin and lysozyme C) across the representations. For more robust training, we performed 5-fold cross-validation for the datasets. Then the trained models were tested on a held-out test set which was 30% of the initial data. Note that, for robustness, we repeated this analysis on 20 distinct random states and reported the mean and standard deviation for the obtained results among all the classification scores.

### EFE Protein Analysis

#### Plasmid Construction

DNA encoding wild-type EFE from strain PK2 was constructed as previously described^59^. Ten EFEs generated by the simple VAE trained on diverse ancestors were chosen randomly for experimental characterization. DNA sequences encoding ancestral and ML-generated EFEs were cloned into the same modified pET28a^+^ vector by IDTdna, with each gene fragment codon optimized for expression in *E. coli* BL21. The modified pET28a^+^ vector includes a sequence encoding an N-terminal His_6_ fused to EFE sequences with a tobacco etch virus (TEV) protease cleavage site, a kanamycin resistance gene, and a T7 promoter regulated by a lac operator sequence to facilitate isopropyl β-D-1-thiogalactopyranoside (IPTG)-induced transcription. Plasmids were transformed into *E. coli* BL21 (DE3) cells (ThermoFisher Scientific, EC0114) following the manufacturer’s instructions. Glycerol stocks were created from single colonies selected for kanamycin resistance (50 mg/mL) on Luria broth (LB) agar plates.

#### Protein Production and Purification

Protein production and purification followed previously published protocols^59^. Briefly, 5-mL LB liquid cultures, supplemented with 50 mg/mL kanamycin, were inoculated from glycerol stocks and grown overnight in a shaking incubator set at 37 °C and 250 rpm. Cultures were diluted into 50 mL terrific broth to an OD_600_ of 0.01-0.1 and shaking was continued at 37 °C and 250 rpm. Once the cells reached an OD_600_ of 0.8-1, they were induced with 0.2 mM IPTG overnight. Cells were pelleted by centrifugation at 6,130 x *g* for 10 min at 4 °C before sonicating them in buffer A (50 mM NaH_2_PO_4_, pH 8.0, containing 500 mM NaCl and 10 mM imidazole) using a Fisherbrand^TM^ 505 sonicator. Samples were sonicated over 10 min at 30% amplitude with 30 s intervals between sonication and ice-chilling steps. Lysates were clarified via centrifugation at 32,280 x *g* for 20 min at 4 °C for subsequent affinity purification via gravity column that was packed with 3 mL of nickel-nitrilotriacetic acid (Ni-NTA) agarose (ThermoFisher Scientific, 88221). Unbound proteins were eluted with 100 mL buffer A. His-tagged EFE protein was eluted in 50 mL of elution buffer (50 mM NaH_2_PO_4_ pH 8.0, containing 500 mM NaCl, and 250 mM imidazole).

Samples were further processed by removing the His-tag with TEV protease. Eluted fractions were first concentrated with Amicon Ultra-15 10 kDa centrifugal filters (EMD Millipore) and buffer exchanged into 50 mM NaH_2_PO_4_ (pH 8.0) containing 500 mM NaCl. The His-tag was subsequently cleaved with TEV protease overnight following manufacturer’s instructions (New England Biolabs, P8112S). The cleaved His-tag fragments were separated from samples with a Ni-NTA column, and the flow-through fractions in buffer A were dialyzed overnight into 25 mM 4-(2-hydroxyethyl)-1-piperazineethanesulfonic acid (HEPES) buffer (pH 8.0) containing 1 mM ethylenediaminetetraacetic acid (EDTA) and 1 mM dithiothreitol (DTT). After adding glycerol to a final concentration of 5% (v/v), samples were flash-frozen in a dry-ice-ethanol bath and stored at -80 °C for future characterization. For both His-tagged and non-His-tagged EFE samples, protein concentrations were determined by analyzing the 280-nm absorbance and using with the calculated molar extinction coefficient (https://web.expasy.org/protparam/). The relative protein sizes and purities were analyzed using 4-20%, pre-cast sodium dodecyl sulfate-polyacrylamide gels (BioRad, #4568096).

#### Protein Thermostability Assay

Protein thermostability was assessed by differential scanning fluorimetry (DSF) for purified proteins following buffer exchange into 50 mM NaH_2_PO_4_ (pH 8.0) and 500 mM NaCl, prior to His-tag cleavage. The Protein Thermal Shift™ Kit from Applied Biosystems™ was used following the manufacturer guidelines. Reactions were prepared in a MicroAmp Fast Optical 96-well plate to a final volume of 20 µL and 0.1 mg/ml protein concentration. Melt curves were acquired over a temperature range of 25 °C to 99 °C using the QuantStudio 5 real-time PCR system. All DSF runs were performed in four technical replicates.

#### EFE Activity Assay

The concentrations of the ethylene product formed during the EFE reaction was determined by gas chromatography as described previously^37^. At room temperature, EFE was added to a final concentration of 250 nM in 2 mL of 25 mM HEPES buffer (pH 7.5) containing 0.5 mM 2OG, 0.5 mM L-Arg, 0.2 mM Fe(NH_4_)_2_(SO_4_)_2_, and 0.4 mM L-ascorbic acid in 10 mm ×16 mm tubes (BD Vacutainer Serum), shaking the tube gently to mix. The reaction was terminated by adding formic acid to a final concentration of 1%. 250 mL of the headspace was withdrawn from the glass tube and injected into a Shimadzu GC-8A gas chromatograph with a flame ionization detector and Porapak N-packed column (80/100 mesh, 2 m × 1/8 in.). Standard curves of ethylene concentrations between 0.8 and 1000 nM were used to calibrate the chromatograph (SCOTTY Analyzed Gases, 99.5%).

## Results

Through both generative and classification tasks, this investigation demonstrated how evolutionary information can be employed in ML platforms to navigate the rugged protein fitness landscape. For generative models, the selection of training data that accurately captures the biological diversity is crucial in improving the quality of the sequences produced. In parallel, the efficacy of fine-tuning language models, a state-of-the-art method for ML predictive tasks, is intrinsically linked to the caliber of the dataset used for refinement.

### IQ-TREE and BAli-Phy-derived ancestral ensembles show similar downstream stability and diversity profiles in the EFE workflow

As part of the initial steps towards curating training datasets enriched with evolutionary information, we evaluated the downstream differences between ancestors inferred by Bayesian inference and maximum likelihood, as implemented in BAli-Phy and IQ-TREE respectively. To improve reconstruction quality, we focused on sampling from high-confidence nodes^60^ in the phylogenetic tree (nodes with SH-aLRT *>* 80% and ultrafast bootstrapping *>* 95%). This approach combines IQ-Tree’s efficiency with a strategy to minimize uncertainty in our ancestral sequence predictions. Across the downstream metrics evaluated here (i.e. sequence variability, semantic diversity, predicted structural confidence, and predicted thermostability) we observed broadly similar profiles for IQ-TREE- and BAli-Phy-derived ancestral datasets. These results do not establish equivalence in ancestral inference quality; rather, they indicate that, for the EFE datasets and downstream ML analyses considered here, IQ-TREE-derived ASR-Dist sequences produced similar stability-and diversity-related outputs at substantially lower computational cost.

### Simple VAE Trained on Resurrected Sequences Generates Novel & Stable EFEs

Three distinct sets of sequences, corresponding to the modern, homogeneous ancestors, and diverse ancestors, were generated. The average atomic distances (RMSDs) between the experimentally determined structure of the wild-type PK2 EFE sequence (PDB ID: 5V2Y) and the AlphaFold2 predicted structures of the training sequences and generated sequences indicate that the generated structures maintained folding patterns similar to the wild-type protein, with the mean RMSD values of 0.43 for modern-derived structures, 0.42 for homogeneous ancestors-derived structures, and 0.39 for diverse ancestors-derived structures. However, ancestral training and ML-generated sequences exhibit greater sequence conservation, as indicated by lower edit distances between training and generated datasets (Figure 3B). The UMAP projection of generated sequences also highlights the broader semantic diversity made possible by implementing ancestral sequences in training (Figure 3C). Despite having more variability in structural quality and sequence composition, modern sequences form tighter clusters in semantic space.

**Figure 3.**
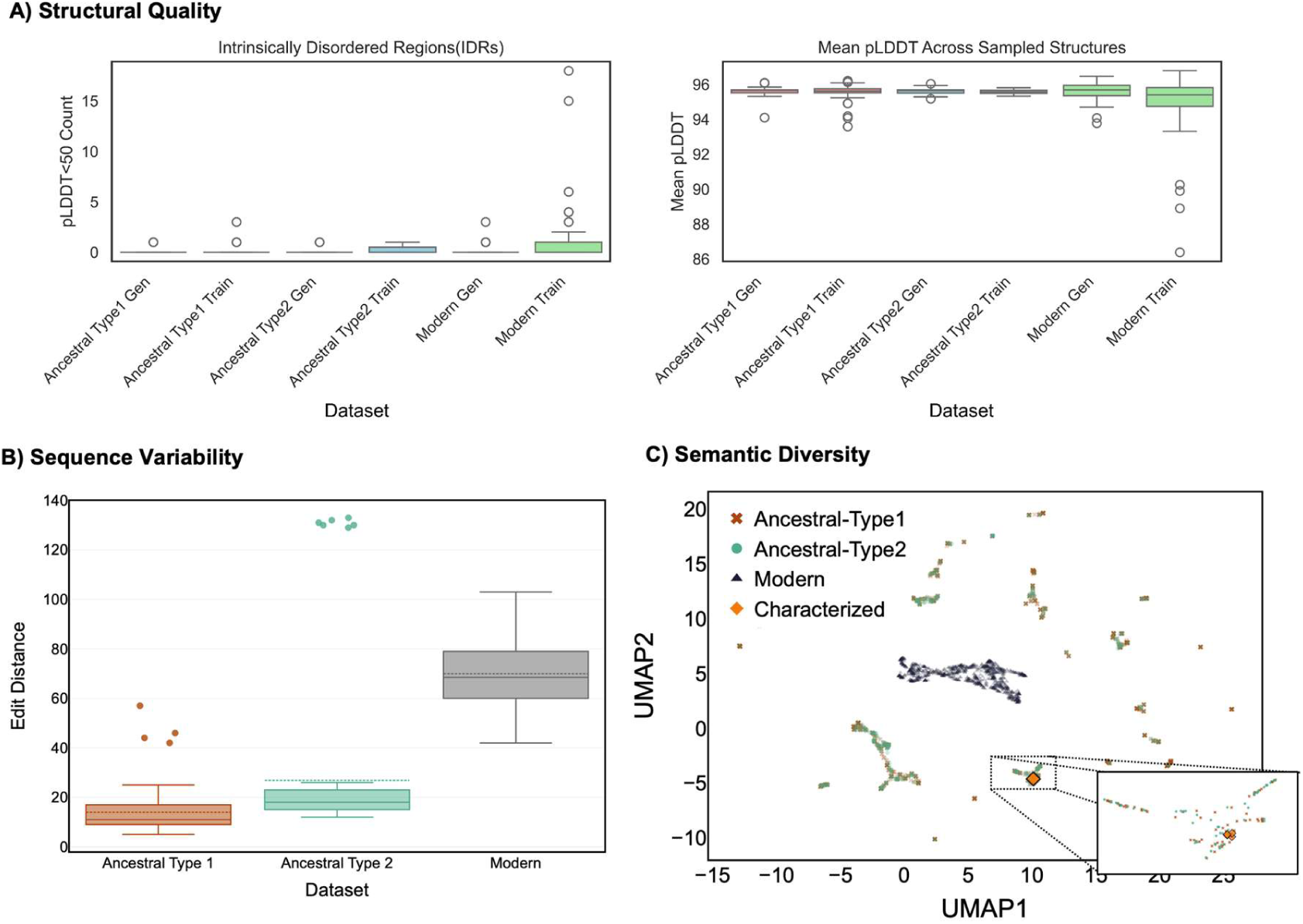
Structural and sequence properties of ancestral and modern EFE sequences before and after generative modeling. **A. Structural Quality Analysis**. Intrinsically Disordered Regions (IDRs) were assessed by the number of amino acid residues where pLDDT<50 and the mean pLDDT scores across sampled structures. Fewer IDRs and higher mean pLDDT scores were generated in ancestral sequences. **B. Sequence Variability Analysis**. Minimum edit distances between training and generated datasets, revealing significantly higher variability in modern sequences (p-value *<* 0.05). **C. Semantic Diversity.** UMAP projection of sampled generated sequences, illustrating distinct clustering patterns with ancestral sequences exhibiting broader distribution. Ancestral sequences are predicted to have improved structural stability, more conserved sequence diversity, and increased semantic diversity compared to modern sequences, while modern sequences show higher sequence variability before and after training.

These results indicate that ASR-enriched training data shifted the VAE-generated sequence distributions toward variants with higher predicted thermostability (Figure 4, Table S1). Both ancestral training datasets and sequences generated from these ancestral training datasets generated sequences from both ancestral types showed no statistically significant difference from their training sets (p-values *>* 0.05), while differing significantly from modern sequences (p-values *<* 10^-8^). Ancestral-based models consistently produced sequences with improved stability compared to modern sequences. Generated sequences closely mirror the stability profiles of their respective training sets. Furthermore, the pLDDT metric suggests improved structural stability for ancestral training and generated sequences with fewer amino acids in intrinsically disordered regions and consistently higher pLDDT scores (Figure 3A). Because these analyses rely on FoldX, AlphaFold-derived confidence metrics, and sequence-embedding projections, we interpret them as computational evidence that ancestral training data bias the model toward stable, well-folded EFE-like sequences, rather than as direct evidence of improved catalytic fitness.

**Figure 4.**
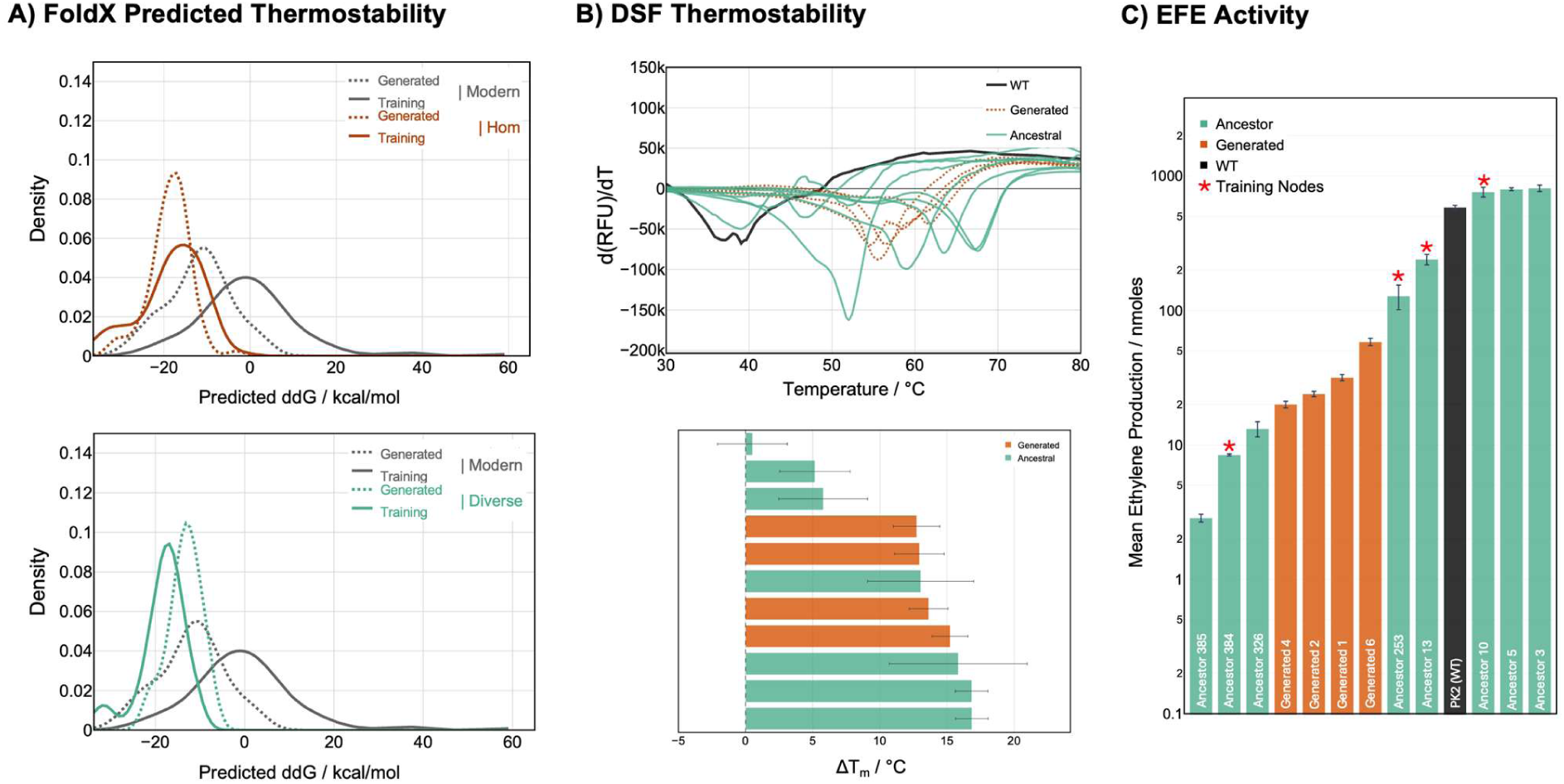
ASR-derived EFE sequences exhibited higher thermal stability compared to modern training sequences and sequences generated by VAE trained on modern sequences. **A.** Predicted stability analysis of homogeneous ancestor sequences and diverse ancestor sequences compared to modern sequences. The density plot shows the thermodynamic stability (ΔΔ*G* values - stability measurement relative to the stability (Δ*G*) of wild-type PK2 EFE) for modern sequences compared to the homogeneous and diverse ancestral protein sequences. **B.** Experimental validation of FoldX-predicted thermostability with differential scanning fluorimetry (DSF). Nearly all the selected ancestral EFE sequences and all the EFE sequences generated from a VAE-trained on ancestral sequences demonstrate significant improvements to thermostability (n=4, p< 0.05). The bar-chart illustrates the ΔTm of ancestors and ML-generated EFE sequences relative to the WT PK2 EFE. **C.** Gas-chromatography confirms EFE activity is conserved. All ancestral EFE sequences and generated EFE produce detectable amounts of ethylene.

### Ancestral and ML-generated EFEs show increased thermostability but ML-generated variants exhibit reduced ethylene-forming activity

The AlphaFold2 and FoldX predictions suggest ancestral and generated EFE sequences have high structural similarity to wild-type PK2 EFE while improving thermostability. To experimentally characterize the stability and activity of these EFE sequences, we randomly selected 4 ML-generated EFE sequences for wet-lab characterization, and 8 ancestral EFE sequences were selected based on UFB and Sh-aLRT confidence values and their relative distance from the reference PK2 EFE. Using DSF to determine T_m_ (i.e., the temperature at which 50% of a given protein sample is unfolded), we found that nearly all of the ancestral sequences (7/8) and all of the ML-generated EFE sequences maintained their folded structures at greater temperatures than the wild-type PK2 EFE (Figure 4B).

When added to a reaction buffer containing 2OG and L-Arg, all ancestral and ML-generated EFE sequences produced detectable amounts of ethylene. Three ancestral EFEs exhibited equivalent or better ethylene formation activity compared to the wild-type PK2 EFE (570 ± 60 nmoles), while the remaining EFE sequences generated between 2.9 nmoles to 240 nmoles of ethylene. Interestingly, despite being generated from an ML model trained on a diverse training set, the ML-generated EFE sequences generated a similar amount of ethylene (between 20 nmoles to 58 nmoles of ethylene). However, this represents a 10-fold decrease in activity compared to the wild-type PK2 EFE. Furthermore, despite the wide range of DT_m_ and ethylene forming activity, there is no meaningful correlation between the two metrics, rs(11)=0.141, p=0.68 (Supplemental Figure S4).

These results suggest that the ancestral-data-trained VAE more readily captured sequence features associated with folding and thermostability than those required for wild-type-like catalytic efficiency. Thus, the generated variants support the utility of ASR-informed training for stability-oriented sequence generation, while also highlighting the need for explicit functional constraints or activity-guided selection in future generative workflows.

### Molecular docking highlights conservation of key protein-ligand interactions in predicted structures of ancestral and generated EFEs

Given the remarkable degree of structural similarity between AlphaFold2-predicted structures of ancestral and generated EFE sequences with the crystal structure of the PK2 EFE (noted above, Figure 5A), we investigated whether protein-ligand interactions were conserved across all generated and ancestral sequences using the joint structural prediction and molecular docking capabilities of AlphaFold3 (AF3). Preliminary testing of AF3 molecular docking with the PK2 EFE sequence demonstrated both high confidence structure prediction (mean pLDDT=95.32) and high structural similarity compared to the previously resolved crystal structure of PK2 EFE (PDB: 5V2Y, RMSD=0.203Å), with all three ligands occupying the same binding pockets and in proximity to the same key binding site residues. When this analysis was expanded to 50 randomly selected ancestral sequences, 50 randomly selected diverse generated sequences, and 50 randomly selected homogeneous generated sequences, several key features of the PK2 EFE structure were observed. All the selected sequences featured a conserved double-stranded beta-helix fold (i.e., jellyroll fold) surrounding the binding site for 2OG, L-Arg, and Mn(II) ion (Figure 5B). The catalytic triad consisting of H189, H268 and D191 (PK2 EFE numbering) common amongst many members of the 2OG/Fe(II)-dependent oxygenase superfamily was present across all predicted structures (Figure 5C).

**Figure 5.**
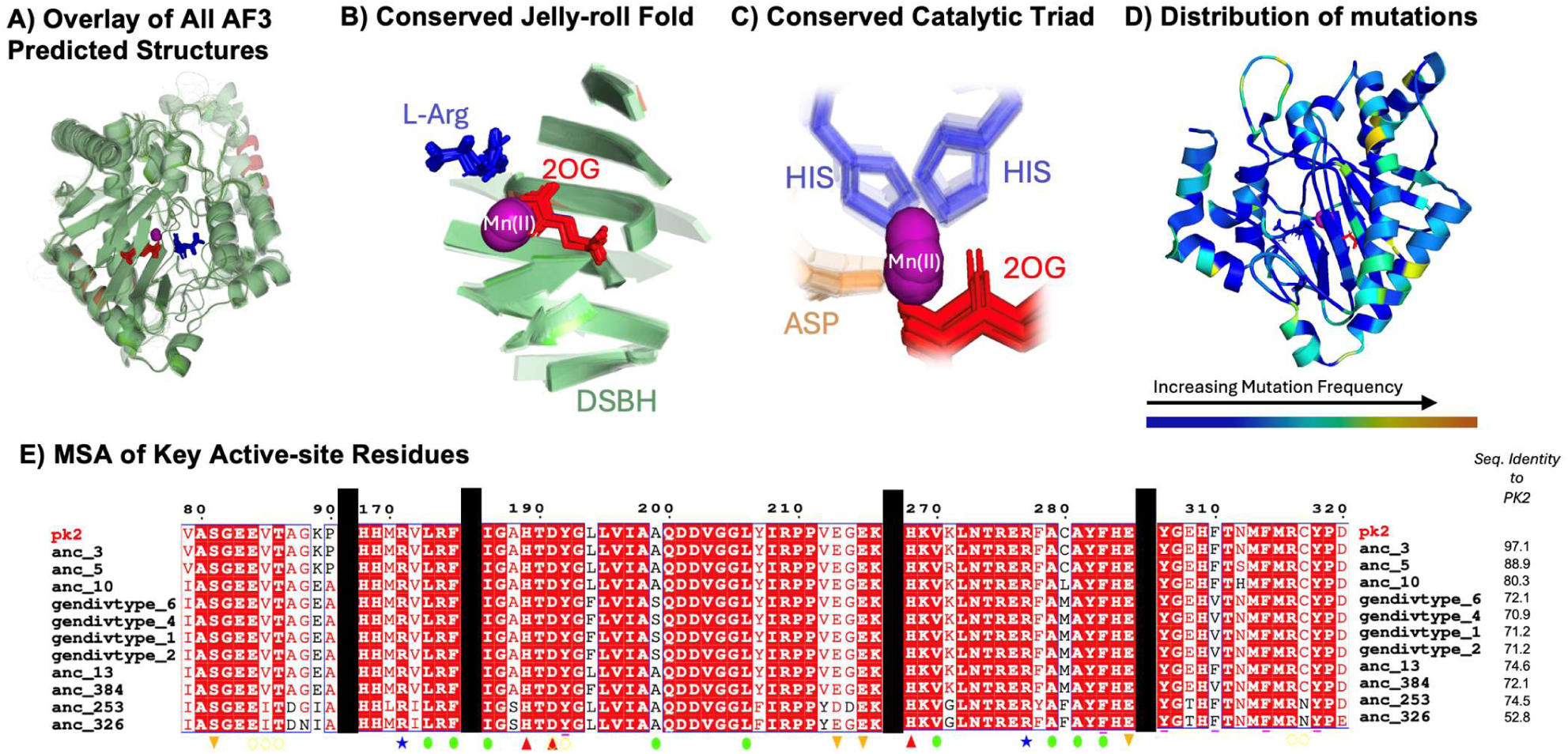
AF3 structural prediction and ligand-placement analyses suggest conservation of key protein-ligand interaction features across ancestral and ML-generated EFEs. A. Overlay of AF3 joint structure predictions and docked complexes for representative ancestral and generated EFE sequences, showing near-identical global folds and highly consistent placement of the three ligands in the active-site region. B) All predicted complexes retain the canonical double-stranded β-helix (jelly-roll) fold that forms the binding pocket for 2-oxoglutarate (2OG), L-arginine (L-Arg), and the divalent metal cofactor (Mn(II)). C) Catalytic triad (PK2 numbering: H189, D191, H268) characteristic of 2OG/Fe(II)-dependent oxygenase, appears preserved across all predicted structures and positioned to coordinate the metal and support catalysis. D) Mapping of mutation frequency onto the protein structure indicates that most substitutions in ancestral and generated sequences are distal to the active site, consistent with preservation of core binding-site geometry and interactions. E) Multiple sequence alignment (MSA) of key active-site positions across wild-type PK2, inferred ancestors, and ML-generated EFEs highlights conservation of residues mediating ligand binding: R171 and R177 (blue stars) support hydrogen bonding to 2OG; L173, F175, I186, L206, V270, A279, A281, and F283 (green ovals) form a hydrophobic environment around 2OG; and E84, T86, D191, Y192, and R316 (yellow open ovals) are conserved at positions contributing to L-Arg interactions. Several non-active-site residues previously implicated in ethylene production (e.g., S81, E213, E215) are also retained, supporting a model in which sequence diversification largely preserves the protein–ligand interaction network required for EFE function. Supplemental Figures S8-S11 show alternative views and labeled positions of panels A-C.

Furthermore, most active site residues were conserved (Figure 5E). These residues include R171 and R177 (blue stars) which participate in hydrogen bonding with 2OG and L173, F175, I186, L206, V270, A279, A281, and F283 (green ovals) which form hydrophobic interactions with 2OG. The hydrogen interaction with L-Arg from E84, T86, D191, Y192, and R316 (yellow empty ovals) were also conserved. Many of the non-active site residues which have been experimentally implicated with ethylene production were also conserved, including S81, E213, E215. However, the bulk of the mutations introduced in ancestral and generated sequences were distal mutations scattered throughout the enzyme (Figure 5D).

### ASR-derived representations are competitive with modern-sequence fine-tuning for stability classification

ASR-derived fine-tuning produced representations that were competitive with InterPro-derived modern-sequence fine-tuning and baseline ESM2 representations across the two protein families tested. The strongest ASR-Dist advantage was observed for KNN classification of lysozyme C, where ASR-Dist achieved modestly higher mean balanced accuracy and ROC-AUC than the InterPro-derived representation. However, across endolysin classifiers and ensemble-based models, performance differences were generally small and often within the observed standard deviations. We therefore interpret these results as evidence that ASR-derived sequence sets may provide useful family-specific representations, rather than as a consistent performance improvement across all classifiers and protein families.

To demonstrate the applicability of this approach for classification tasks, we fine-tuned the ESM2 model on three distinct datasets to obtain protein representations termed Inter-Pro, ASR-Max, and ASR-Dist. Intriguingly, the representations derived from fine-tuning ESM2 with ancestral data exhibited comparable or enhanced performance relative to those derived from modern data in determining if a given sequence is stable (ΔΔ*G <* −0.5 *kcal/mol*) or unstable (ΔΔ*G >* 0.5 *kcal/mol*) for endolysin (Table 2, Table S2) and lysozyme C (Table 3, Table S3) proteins. Our ASR-Dist method, which leverages uncertainty in ASR, has shown improved performance over other representations for KNN classification. However, it demonstrated comparable or improved performance over InterPro-derived representations for ensemble-based classifiers (i.e., Random Forest and XGBoost).

**Table 2.**
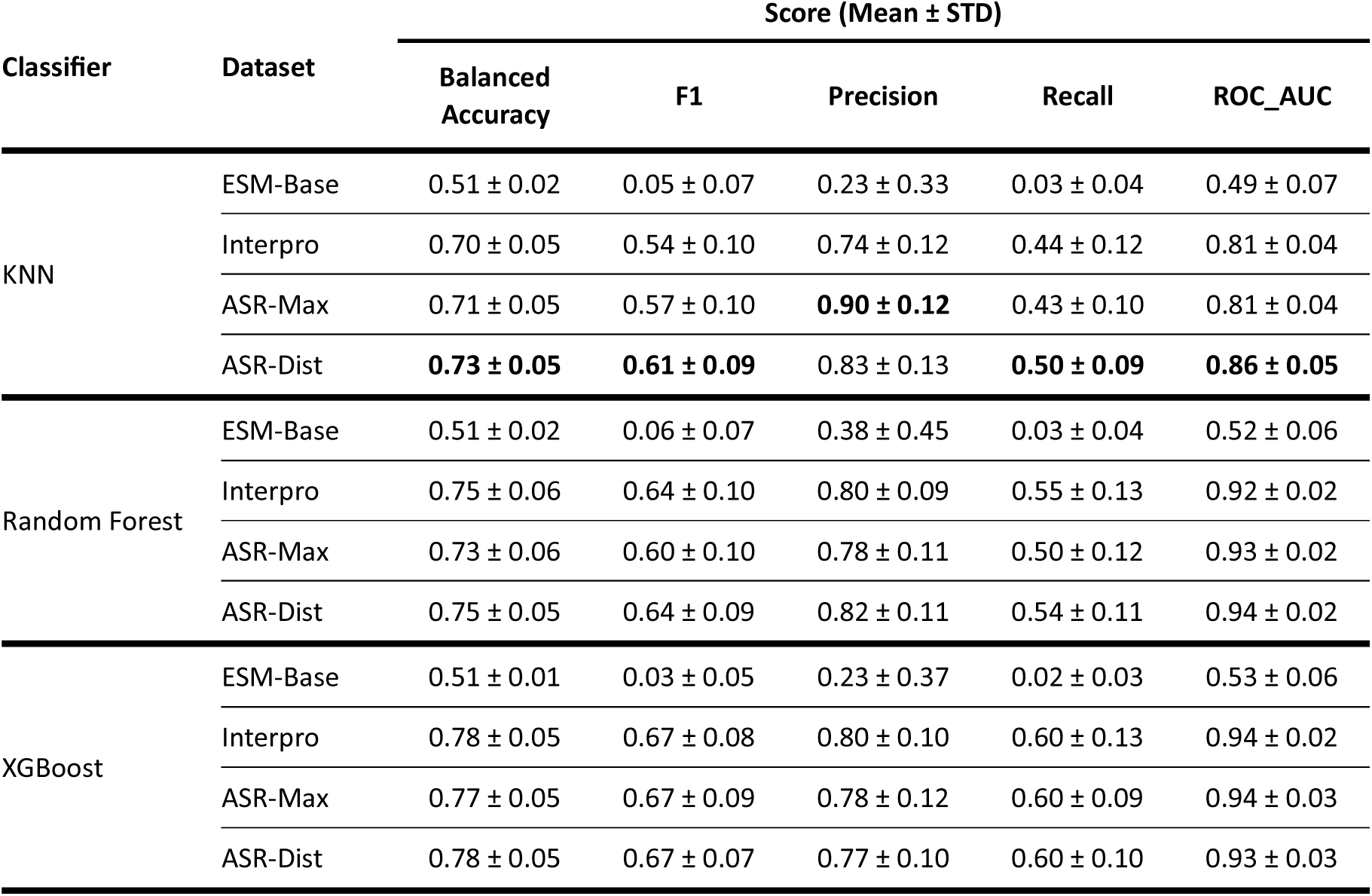
Comparison of Classifier Performance Across Fine-Tuned Representations – Lysozyme C.

**Table 3.**
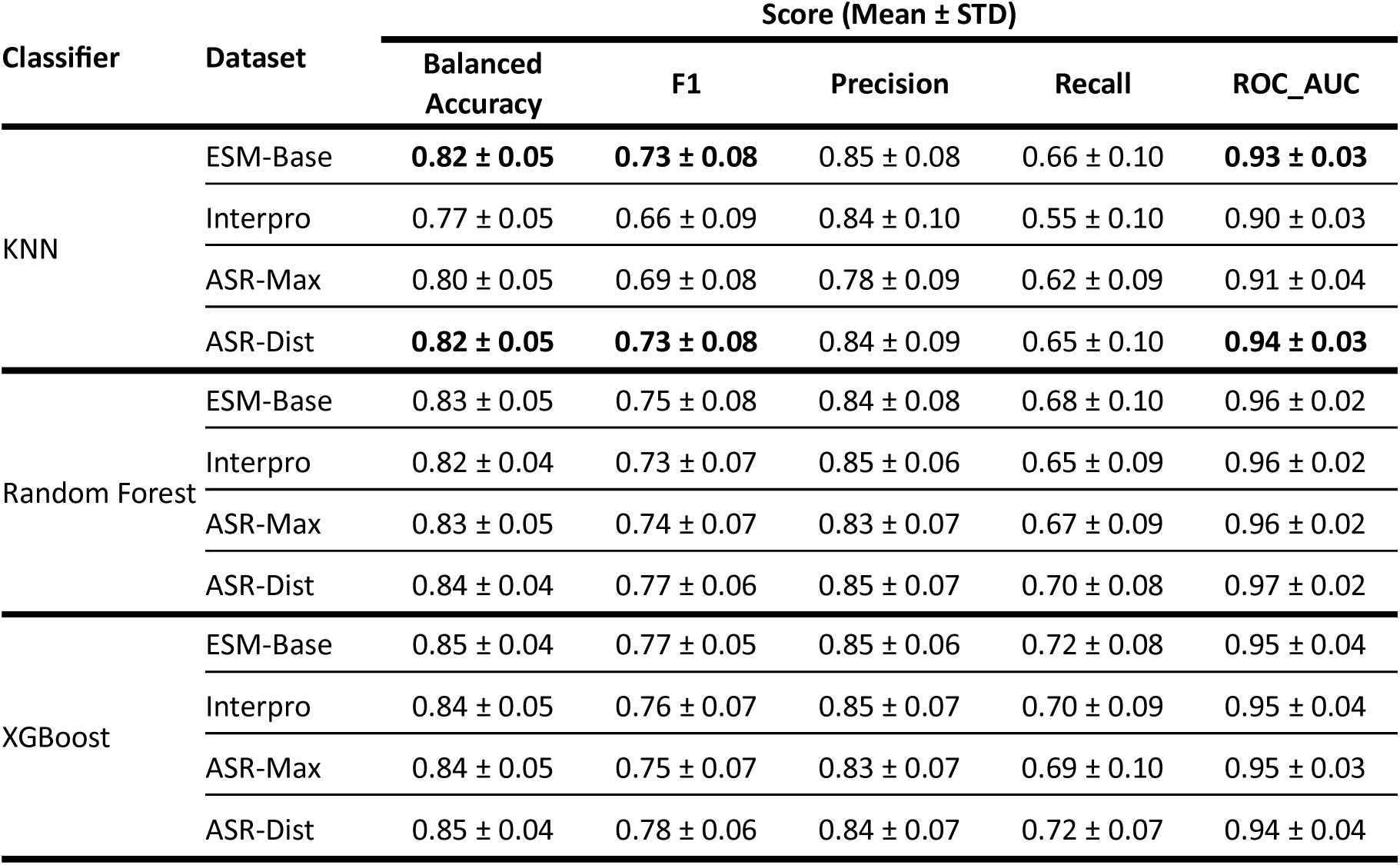
Comparison of Classifier Performance Across Fine-Tuned Representations – Endolysin.

## DISCUSSION

In this work, we explored the integration of evolutionary information into ML models for protein engineering. Specifically, we evaluated whether ASR-derived sequence ensembles can serve as data augmentation sets for ML-guided protein engineering. In the EFE generative-modeling workflow, computational and wet-lab studies showed that ancestral and near-ancestral training datasets shifted VAE-generated sequences toward improved thermostability compared to VAE-generated sequences trained on modern sequences alone. However, the ML-generated EFEs exhibited lower ethylene-forming activity than wild-type EFE, indicating that the model captured stability-associated sequence features more effectively than catalytic optimization. In the ESM2 fine-tuning workflow, ASR-derived representations were competitive with modern-sequence fine-tuning for stability classification, with modest advantages in selected classifier/family combinations but no universal improvement across all models tested.

Our approach combined ASR with generative modeling and language model fine-tuning to enhance both sequence generation and fitness prediction. The results demonstrated that sequences generated from evolutionary-informed datasets exhibited improved characteristics, particularly in thermal stability, compared to those derived from models trained solely on modern sequences. This highlights the potential of ASR to enrich training data for generative models, leading to more diverse and higher-quality outputs.

Briefly comparing maximum likelihood and Bayesian inference approaches to infer ancestral training sequences, IQ-TREE and BAli-Phy workflows produced ancestral datasets with similar downstream stability and diversity profiles in this EFE case study. This finding should not be interpreted as demonstrating equivalent ancestral inference quality. Rather, it suggests that for the downstream ML objectives examined here, IQ-TREE-derived marginal posterior sampling was sufficient to generate sequence sets with similar predicted stability and diversity properties, while being substantially more computationally tractable.

A key factor potentially contributing to this similarity is the sampling strategy. By incorporating sequences from all nodes in the phylogenetic tree, both methods likely capture a wide range of ancestral states, including those with varying degrees of certainty. This comprehensive sampling may balance out the theoretical advantages of each approach, resulting in similar overall performance. The stability results indicate that both methods generate sequences with comparable thermodynamic properties, suggesting IQ-Tree’s point estimates are as robust as BAli-Phy’s distribution-based estimates for maintaining protein stability. The equivalent sequence variability and semantic diversity imply that IQ-Tree’s maximum likelihood approach explores sequence space as effectively as BAli-Phy’s Bayesian method.

As demonstrated in both computational predictions and wet-lab characterization, the ancestral training sequences and evolutionary-informed generated sequences exhibit enhanced thermostability. The precise mechanisms underlying this shift remain unclear, it is possible that the generative model, when applied to modern sequences, mitigates noise from low-quality sequences (in terms of epistasis and folding). Conversely, the ancestral training data may inherently encompass stabilizing interactions. Consequently, the generated sequences from ancestral proteins consistently demonstrate superior stability compared to those derived from modern sequences.

These findings highlight the potential benefits of incorporating evolutionary data in generative models for protein design. Ancestral sequences appear to combine desirable properties such as structural stability with an expanded exploration of sequence space. This combination could prove valuable in generating novel protein variants that maintain crucial ancestral characteristics while introducing functional innovations. By leveraging the broader semantic diversity of ancestral sequences, generative models could potentially access a wider range of protein designs, opening new avenues for engineering proteins with enhanced or altered functions while preserving their fundamental structural integrity.

While our study provided valuable insights, it’s important to note that we only scratched the surface of the vast sequence space available through ASR. For instance, considering the 186 high-quality nodes from our lysozyme ancestral tree reconstruction, with 15 variable positions and 4 possible amino acids per position, the theoretical sequence space encompasses approximately 200 billion unique sequences. Our approach, utilizing clustering algorithms and strategic sampling methods, aimed to capture a representative subset of this diversity. Despite these efforts to efficiently sample the sequence space, future work could focus on developing more sophisticated sampling techniques to better explore the available sequence space which may reveal even more beneficial protein variants.

To evaluate the utility of ancestral sequences on downstream stability prediction tasks, we fine-tuned the evolutionary scale ESM protein language model with reconstructed ancestral data to obtain evolutionary-driven protein representations for endolysin and lysozyme C families. For lysozyme C, ancestral-based representations showed promising results, outperforming the baseline ESM in KNN classification and matching the established InterPro method for other classification models. For endolysin, our novel ASR-Dist method performed comparably to the baseline and other fine-tuning approaches across various classification metrics. While not consistently outperforming existing methods, ASR-Dist demonstrated stable performance across both simple and complex classification models, suggesting the potential for use in this data-centric approach for enhancing protein representations.

This study advances protein engineering by integrating ASR into generative modeling and representation learning. It demonstrates the generation of sequences with ancestral-like properties using machine learning. While previous work such as local ancestral sequence embedding (LASE) introduced ASR in representation learning using transformers trained from scratch, this approach explores fine-tuning existing models with various evolutionary and modern sequence datasets, addressing potential over-fitting issues. Of particular interest for protein engineering, the generative modelling aspect of this study recapitulates prior work leveraging the relative thermostability of ancestral sequences for protein engineering tasks and expands this principle through more nuanced sampling approaches of generated sequences. This comprehensive incorporation of ASR-driven protein sequences and their impact on both generative modeling and representation learning provides a nuanced view of evolutionary information’s role in protein engineering.

### Limitations of the study

While this study illuminates the potential of integrating evolutionary data into ML models, it also highlights the necessity for further investigation. We computationally validated the stability and diversity of the sequences generated for the generative task, but experimental validation is crucial to understanding the functional advantages that evolutionary signals may confer.

## Supplemental information index

Figure S1. Phylogenetic Tree of EFE

Figure S2. Overview of how near-ancestors are constructed using the maximum likelihood approach

Figure S3. Overview of variational autoencoder (VAE) architecture used for generating novel protein sequences

Figure S4. Correlation analysis between thermostability and ethylene production

Figure S5. Sequence Identity Matrix

Figure S6. MSA of experimentally characterized EFE-like sequences with secondary structure annotated

Figure S7. Amino acid diversity among variants

Figure S8. Protein Ligand Interaction Features (protein level)

Figure S9. AF3 structural prediction and docking

Figure S10. Protein Ligand Interaction Features (2OG binding pocket)

Figure S11. Protein Ligand Interaction Features (L-Arg binding pocket)

Figure S12. Protein-ligand interaction profile comparison

Table S1. Statistical Analysis of Generated Protein Stability

Table S2. Statistical Analysis of Protein Representation Performance – Endolysin

Table S3. Statistical Analysis of Protein Representation Performance - Lysozyme C

Supplemental Note 1 – Description and Summary of Generative VAE Architecture

Supplemental Methods – BAli-Phy replicate workflow, ASR generation, and AP-LASR integration

## Acknowledgments

This work was funded by the USDA (NIFA-AFRI: grant 13700968 to D. R. W.), the National Science Foundation (grant 2203472 to R.P.H.), and the Department of Chemical Engineering and Materials Science at Michigan State University. The support and critical analysis by all member of the Woldring Lab were critical in developing this report.

## Author contributions

Conceptualization, M.M., P.F., and D.R.W.; methodology, M.M., N.P., P.F., and D.R.W.; investigation, M.M., N.P., and D.R.W.; writing – original draft, M.M., N.P., and D.R.W.; writing – review & editing, M.M., N.P., P.F., A.M., R.P.H. and D.R.W.; funding acquisition, D.R.W. and R.P.H.; resources, D.R.W.; supervision, D.R.W.

## Declaration of generative AI and AI-assisted technologies

During the preparation of this work, the author(s) used ChatGPT to improve writing quality for select passages. After using this tool or service, the author(s) reviewed and edited the content as needed and take(s) full responsibility for the content of the publication.

## Resource availability

### Lead contact

Requests for further information and resources should be directed to and will be fulfilled by the lead contact, Daniel Woldring.

## Materials availability

This study did not generate new materials.

## Data and code availability

- All original code is publicly available and can be obtained at GitHub: https://github.com/WoldringLabMSU/Evo-Seq.
- All original data, training sequences, and generated sequences are publicly available at 10.5281/zenodo.18394294.
- Any additional information required to reanalyze the data reported in this paper is available from the lead contact upon request.
- Specific Datasets:
- A step-by-step protocol for running the BAli-Phy replicate workflow on SLURM (including generation of MAP.tree, consensus alignments, and ASR.state) and the associated scripts are provided in the Supplementary Information and in the GitHub repository.

