## Supplemental Figures and Tables for "EvoSeq-ML: Advancing Data-Centric Machine Learning with Evolutionary-Informed Protein Sequence Representation and Generation"

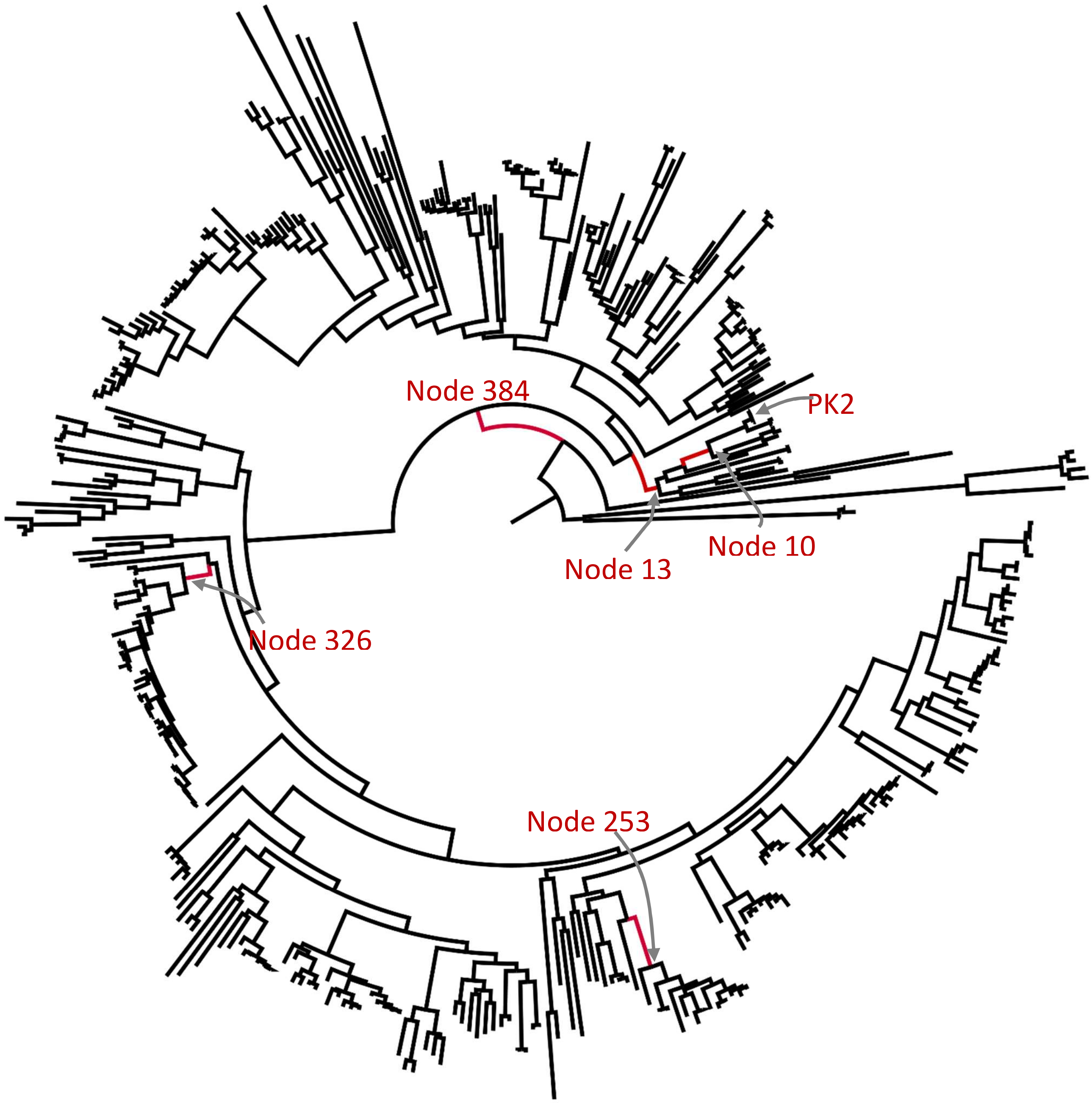

**Supplemental Figure S1.** Phylogenetic Tree of PK2/EFE proteins.

Nodes of interest (e.g. those experimentally characterized and those associated with near-ancestor training data for generative models) are highlighted in red. Raw treefile data available: https://github.com/WoldringLabMSU/Evo-Seq

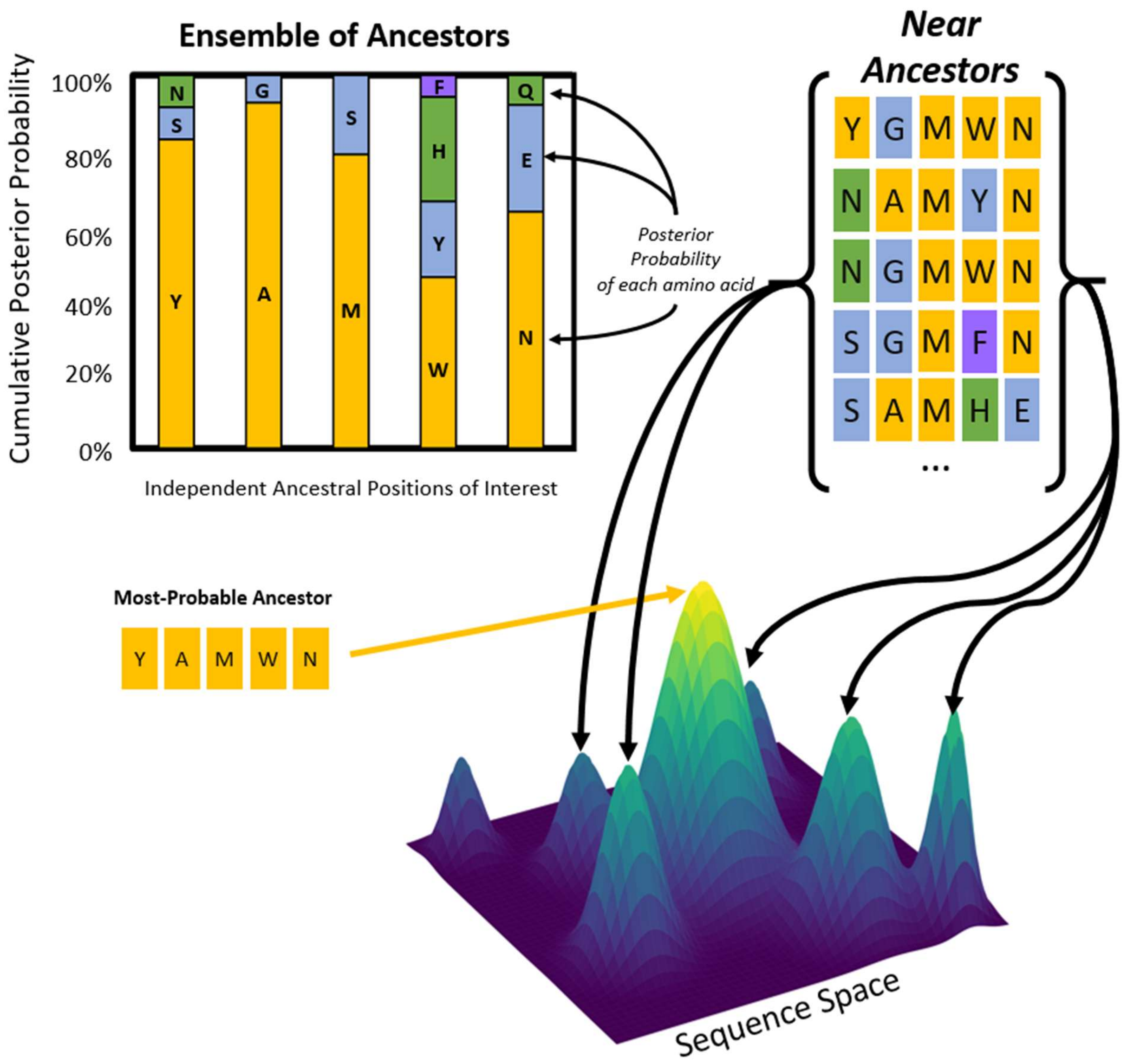

**Supplemental Figure S2**: Overview of how near-ancestors are constructed using the maximum likelihood approach (e.g. IQ-Tree).

For any given node on a phylogenetic tree, a consensus ancestor is inferred based on the most probable amino acid at each position of protein sequence. A subset of these positions have multiple amino acids that are plausible. Therefore, by sampling from the plausible amino acids at each position, an ensemble of ancestors (near-ancestors) can be produced from a single node within a phylogenetic tree.

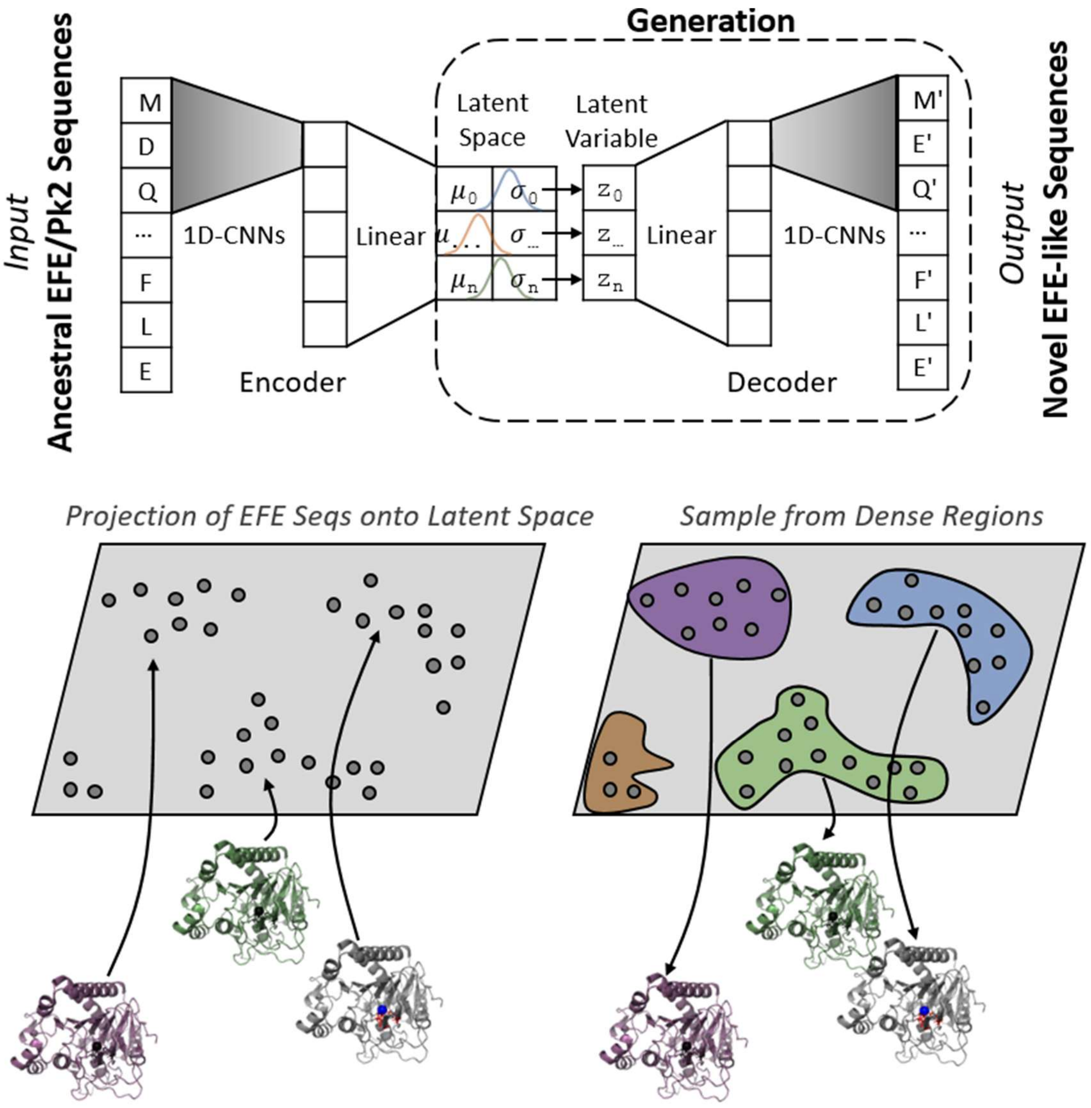

**Supplemental Figure S3**: Overview of variational autoencoder (VAE) architecture used for generating novel protein sequences.

Evo-Seq's generative model is a VAE that learns to compress an aligned protein sequence into a compact numeric code and then reconstruct it. A collection of protein sequences (e.g. ensembles of near-ancestors produced by either maximum likelihood or Bayesian inference) are "one-hot encoding" plus a 1D CNN feature extractor. **Encoder:** each sequence is first represented as a fixed-length, position-by-position one-hot tensor over an alphabet of 20 amino acids plus a gap character; a 1D convolutional layer (with batch normalization and ReLU) scans along the sequence to extract local patterns, and fully connected layers compress these features into two vectors: a latent mean and latent log-variance. **Latent space:** the model samples a continuous latent vector z using the reparameterization trick and is regularized by a KL-divergence term that encourages these latents to follow a standard normal distribution N(0,I), creating a smooth space where nearby points tend to decode to similar sequences. **Decoder:** a mirrored stack of fully connected layers expands z back to sequence-length features, followed by a transposed 1D convolution that outputs a per-position score/probability for each amino acid (and gap); the reconstructed sequence is obtained by choosing (or sampling) an amino acid at each position from these outputs.

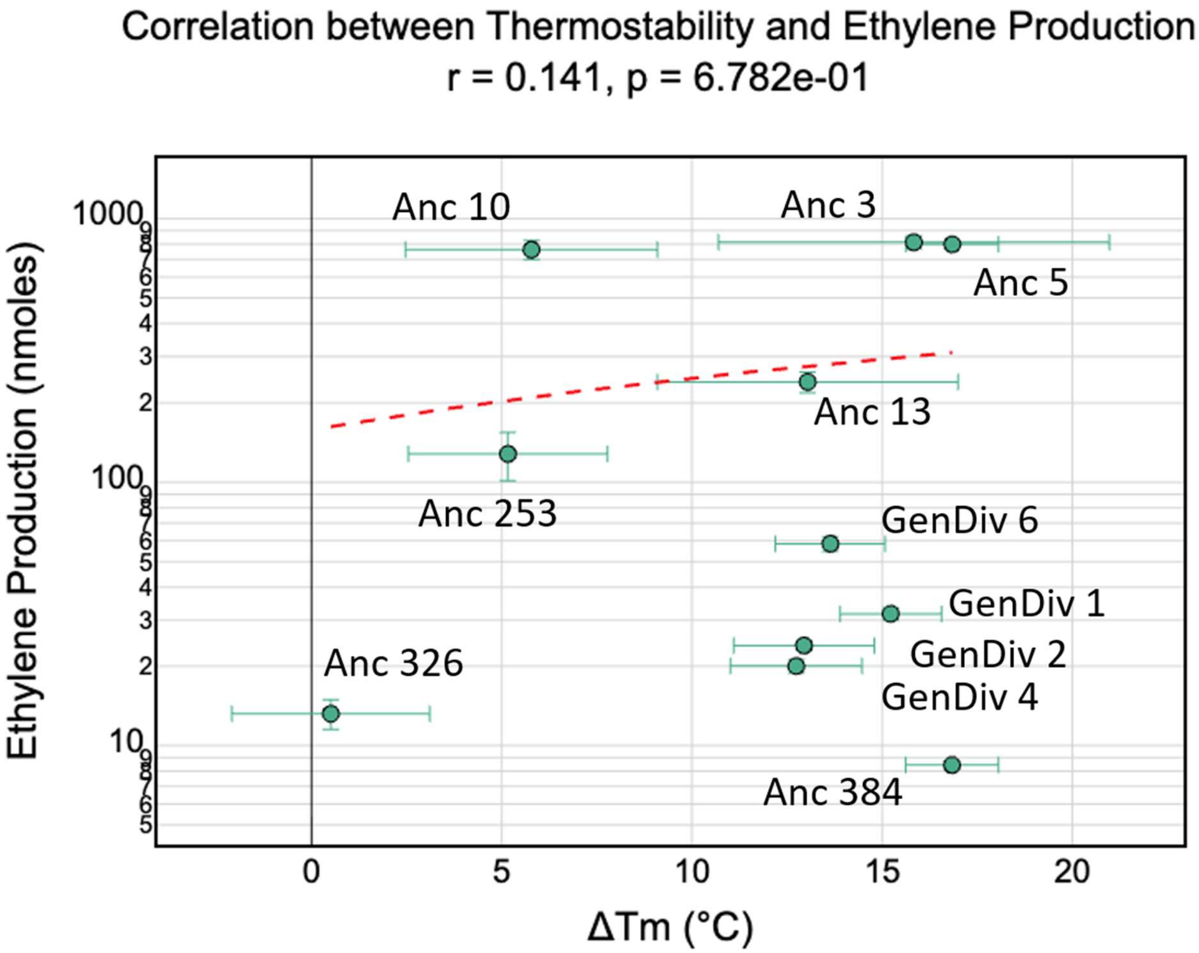

**Supplemental Figure S4**: Correlation analysis between thermostability and ethylene production

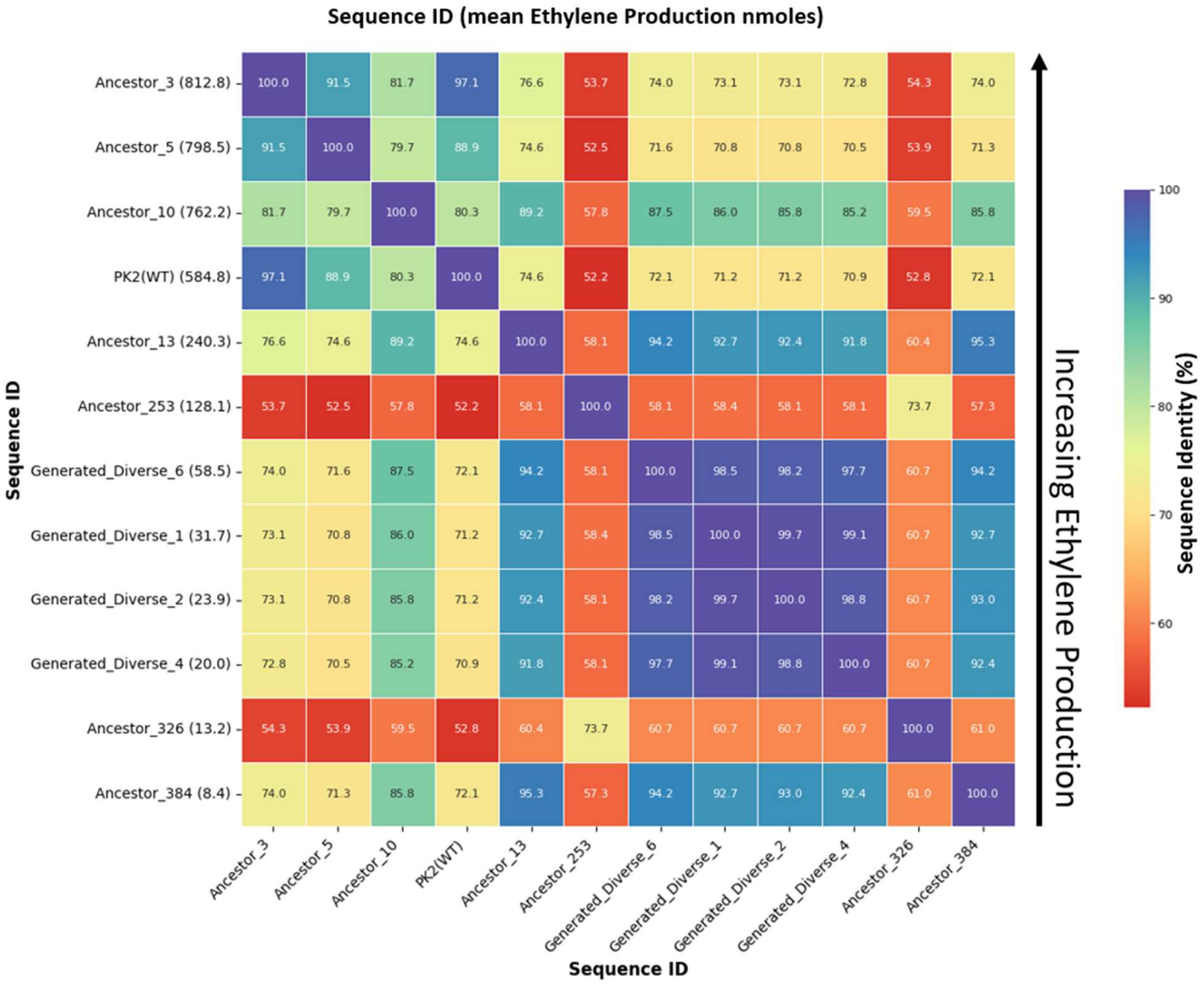

**Supplemental Figure S5**: Sequence Identity Matrix.

The heatmap indicates the percentage of amino acid similarity between each pair of protein sequences. The rows are arranged in descending order based on each variant's ethylene production (values shown in nmoles within parentheses along left side of heat map).

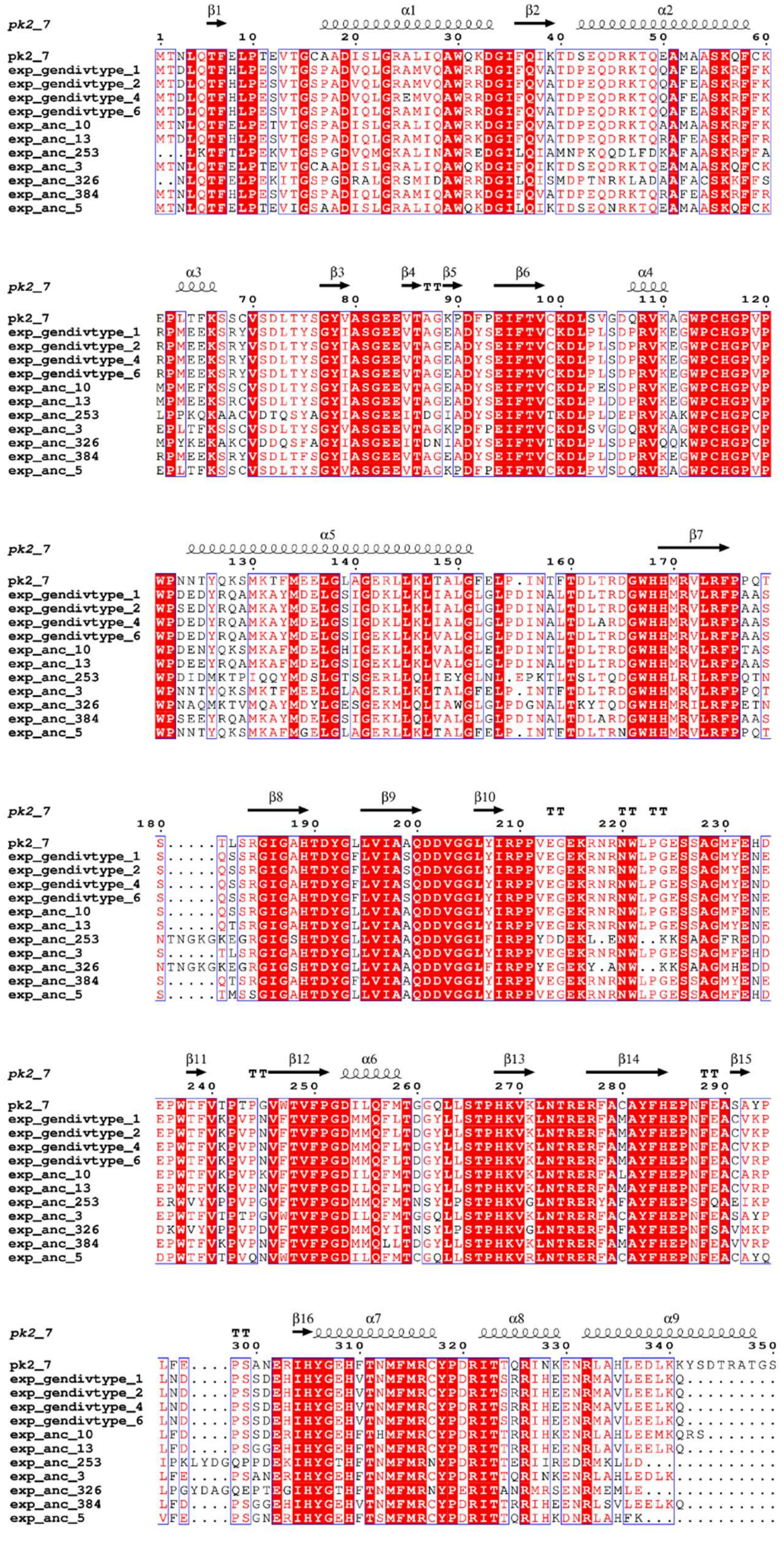

**Supplemental Figure S6**: MSA of experimentally characterized EFE-like sequences with secondary structure annotated.

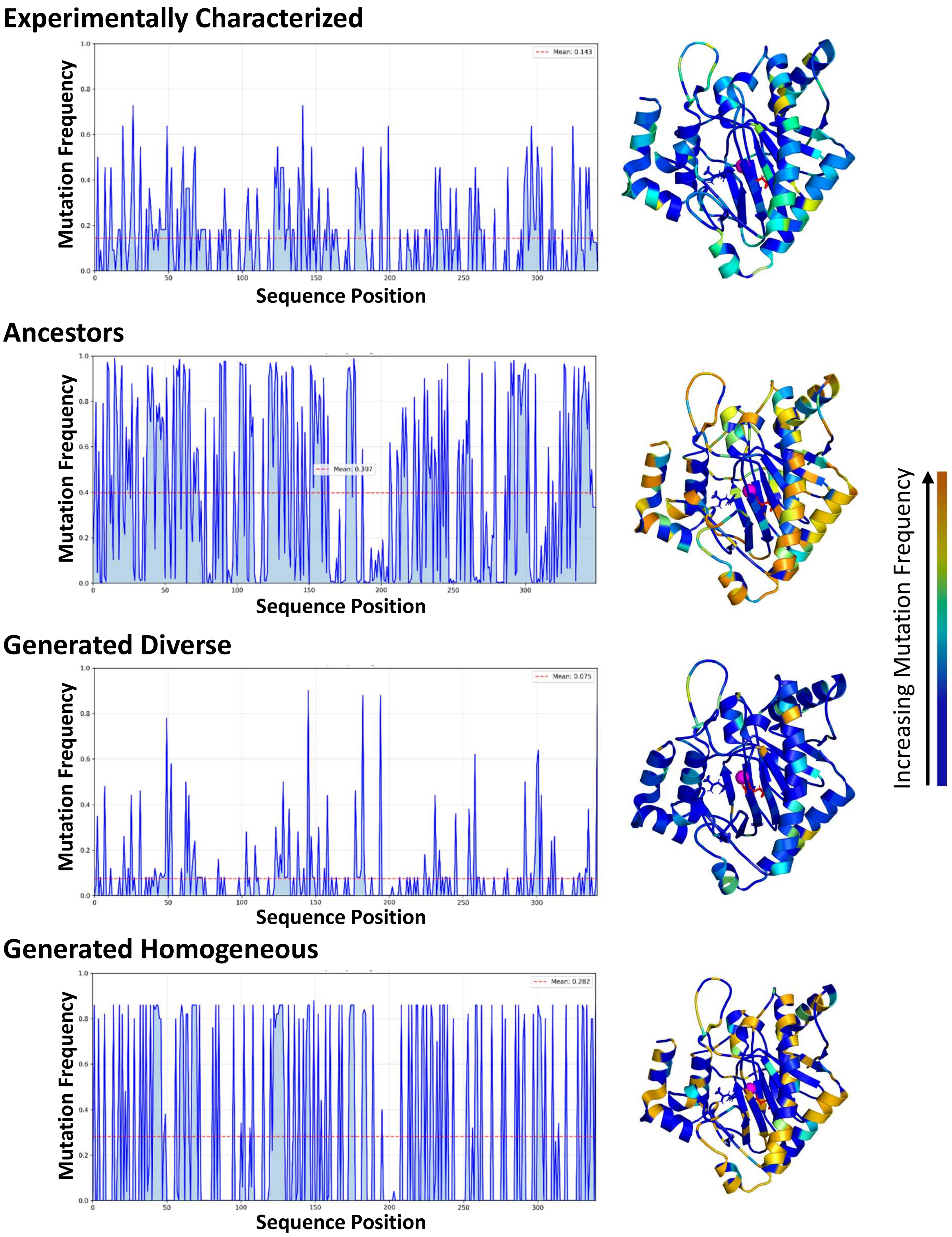

**Supplemental Figure S7:** Amino acid diversity among variants at individual positions within either the subset of experimentally characterized EFE-like variants, inferred ancestral EFEs, or the VAE generated variants.

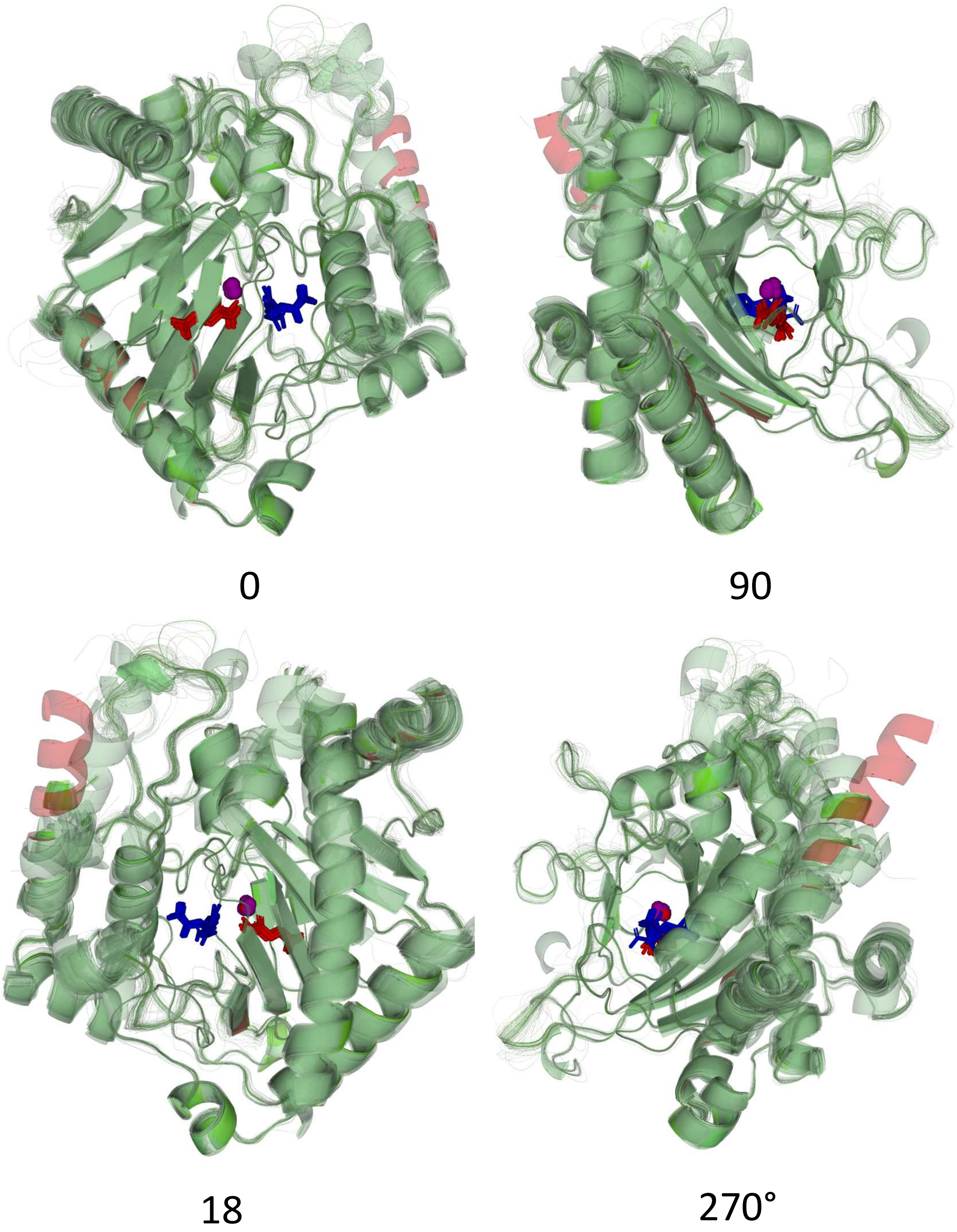

**Supplemental Figure S8**:Protein Ligand Interaction Features (protein level). AF3 structural prediction and docking reveal conserved protein-ligand interactions across ancestral and ML-generated EFEs.

Here we show overlays of AF3 joint structure predictions and docked complexes for representative ancestral and generated EFE sequences, showing near-identical global folds and highly consistent placement of the three ligands in the active-site region.

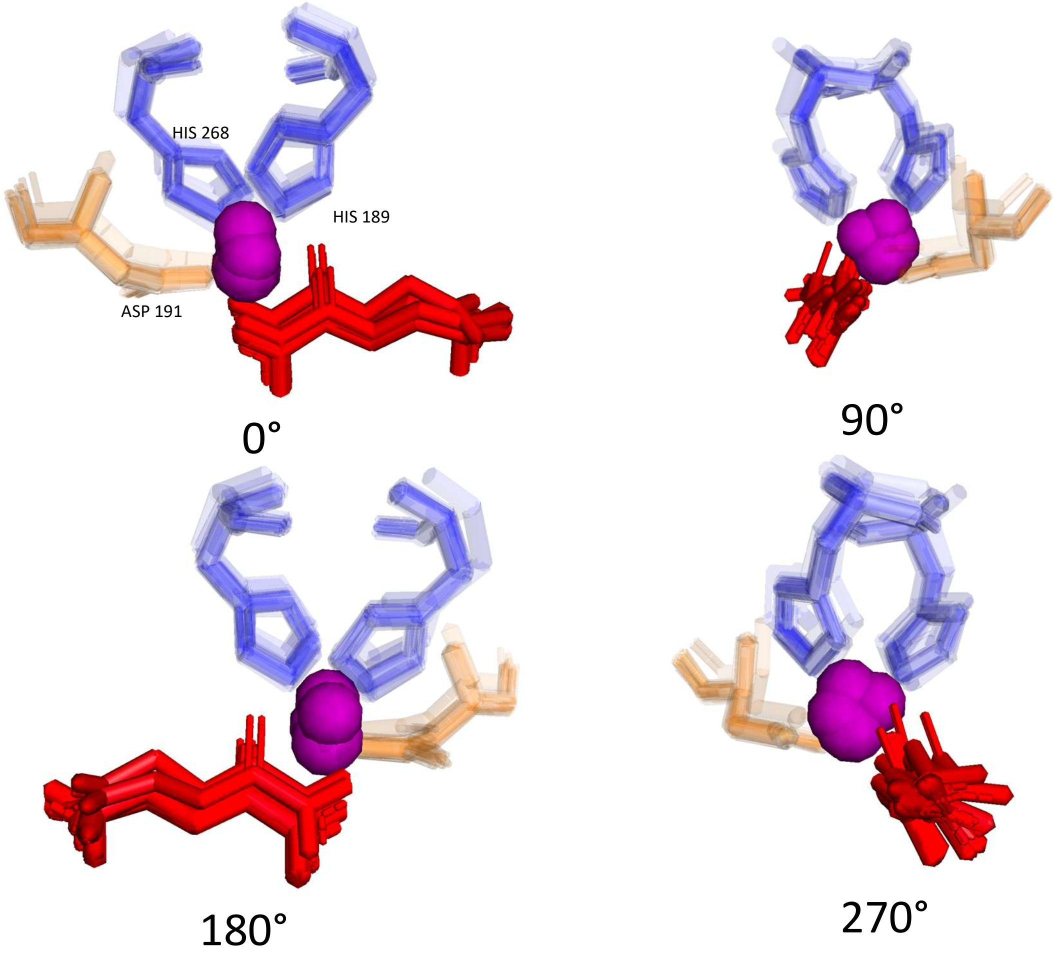

**Supplemental Figure S9:** AF3 structural prediction and docking reveal conserved protein-ligand interactions across ancestral and ML-generated EFEs.

Here we show a close-up of the conserved catalytic triad (PK2 numbering: H189, D191, H268) characteristic of 2OG/Fe(II)-dependent oxygenases, preserved across all predicted structures and positioned to coordinate the metal and support catalysis.

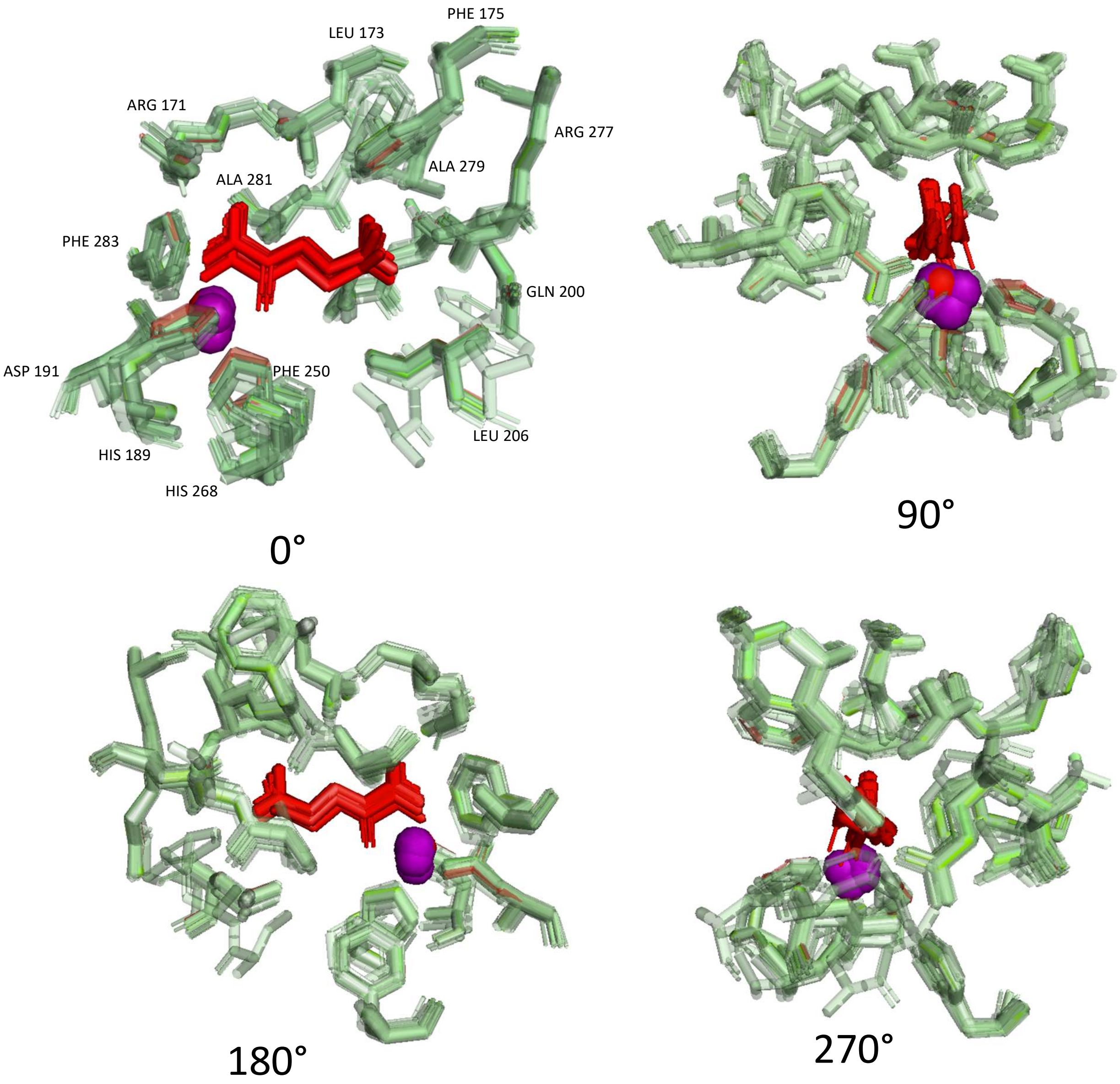

**Supplemental Figure S10**: Protein Ligand Interaction Features (2OG binding pocket).

AF3 structural prediction and docking reveal conserved protein-ligand interactions across ancestral and ML-generated EFEs. Here we show superimposed structures of ancestral and generated EFEs with residues proximal to the canonical double-stranded β-helix (jelly-roll) fold surrounding 2-oxoglutarate (**2OG**) and the divalent metal cofactor (**Mn(II)**).

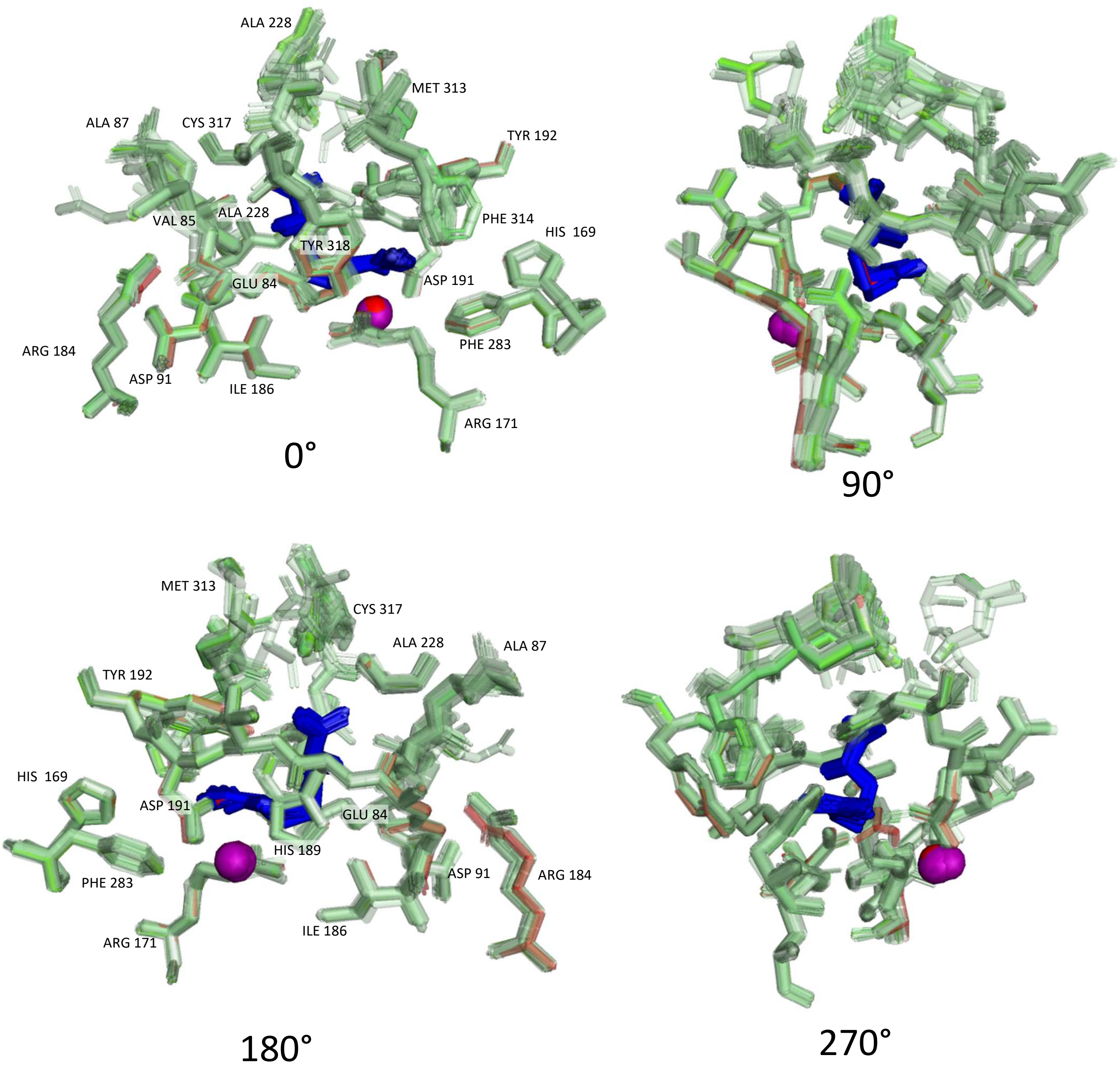

**Supplemental Figure S11:** Protein Ligand Interaction Features (L-Arg binding pocket).

AF3 structural prediction and docking reveal conserved protein-ligand interactions across ancestral and ML-generated EFEs. Here we show superimposed structures of ancestral and generated EFEs with residues proximal to the canonical double-stranded β-helix (jelly-roll) fold surrounding L-arginine (**L-Arg**) and the divalent metal cofactor (**Mn(II)**).

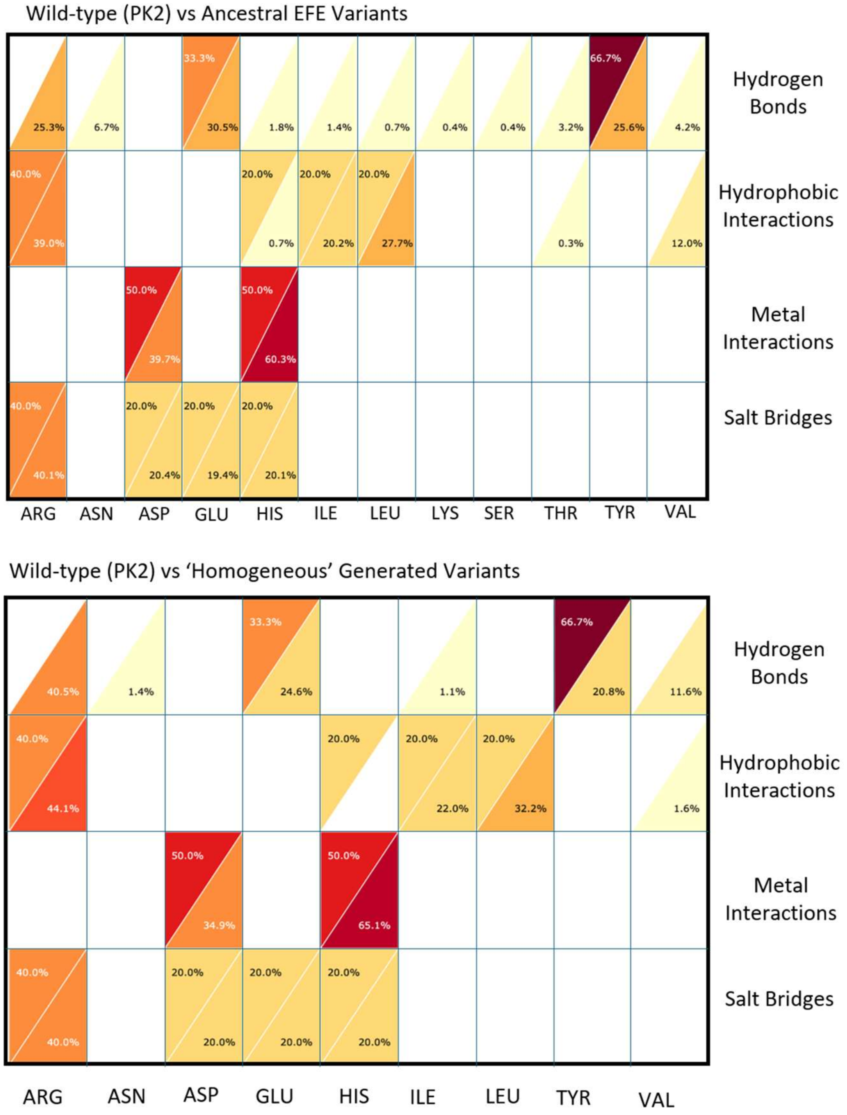

**Supplemental Figure S12:** Protein-ligand interaction profile comparison between wild-type EFE PK2 and either ancestral EFEs (top) or ‘homogeneous’ generated sequences (bottom).

Collections of 50 randomly selected ancestral sequences, and 50 randomly selected “homogeneous” generated sequences were structurally modeled and co-folded with their substrates using AlphaFold3 to investigate potential differences in protein-ligand interactions with the wild-type EFE bound to Mn(II), 2-oxoglutarate (2OG), and L-arginine (L-Arg). The resulting structures were analyzed with the protein-ligand interaction profiler (PLIP) to index potential protein-ligand interactions. The upper-left triangle portions in each cell correspond to the wild-type EFE PK2 protein-ligand interaction profile; whereas the bottom-right triangle portion of the cells in the top table correspond to the ancestral EFE variants, and the bottom-right triangle portions of the cells in the bottom table correspond to the ‘homogeneous’ generated variants. The values shown within a single triangle reflect the relative frequency of a particular amino acid participating in one type of intermolecular interaction. For example, 66.7% of hydrogen-bonding between wild-type EFE and 2OG/L-Arg ligands include a tyrosine (TYR); however, among ancestral EFE variants, only 25.6% of H-bonds with 2OG/L-Arg ligands included a tyrosine, and 20.8% of H-bonds among ‘homogeneous’ generated EFE variants involved tyrosine.

**Supplemental Table S1**: Statistical Analysis of Generated Protein Stability.

Analysis used Welch two-sample t-tests while applying Holm multiple-comparisons correction.

| | Modern-Generated | Ancestral-Type1-Generated | Ancestral-Type2-Generated | Modern-Training | Ancestral-Type1-Training | Ancestral-Type2-Training |
|---|---|---|---|---|---|---|
| Modern-Generated | 1 | 5.40E-10 | 1.39E-08 | 1.21E-08 | 2.18E-07 | 6.66E-06 |
| Ancestral-Type1-Generated | 5.40E-10 | 1 | 0.6972 | 2.31E-31 | 0.2143 | 0.0608 |
| Ancestral-Type2-Generated | 1.39E-08 | 0.6972 | 1 | 3.61E-28 | 0.4168 | 0.1519 |
| Modern-Training | 1.21E-08 | 2.31E-31 | 3.61E-28 | 1 | 1.65E-27 | 2.59E-24 |
| Ancestral-Type1-Training | 2.18E-07 | 0.2143 | 0.4168 | 1.65E-27 | 1 | 0.5069 |
| Ancestral-Type2-Training | 6.66E-06 | 0.0608 | 0.1519 | 2.59E-24 | 0.5069 | 1 |

**Supplemental Table S2**: Statistical Analysis of Protein Representation Performance – Endolysin.

Dunn Test for Statistical Significance after Kruskal-Wallis Test for Different Metrics.

| Group1 | Group2 | P-Value | Classifier | Metric |
|---|---|---|---|---|
| InterPro | ASR-Dist | 0.002346 | KNN | ROC_AUC |
| ASR-Dist | InterPro | 0.002346 | KNN | ROC_AUC |
| InterPro | ASR-Dist | 0.011602 | KNN | Recall |
| ASR-Dist | InterPro | 0.011602 | KNN | Recall |
| InterPro | ASR-Dist | 0.013327 | KNN | Balanced_Accuracy |
| ASR-Dist | InterPro | 0.013327 | KNN | Balanced_Accuracy |
| ASR-Max | ASR-Dist | 0.024595 | KNN | ROC_AUC |
| ASR-Dist | ASR-Max | 0.024595 | KNN | ROC_AUC |
| InterPro | ASR-Dist | 0.042191 | KNN | F1 |
| ASR-Dist | InterPro | 0.042191 | KNN | F1 |
| InterPro | ASR-Max | 0.113910 | KNN | Recall |
| ASR-Max | InterPro | 0.113910 | KNN | Recall |
| InterPro | ASR-Max | 0.214170 | KNN | Balanced_Accuracy |
| ASR-Max | InterPro | 0.214170 | KNN | Balanced_Accuracy |
| ASR-Max | ASR-Dist | 0.557544 | KNN | F1 |
| ASR-Dist | ASR-Max | 0.557544 | KNN | F1 |
| InterPro | ASR-Max | 0.772055 | KNN | F1 |
| ASR-Max | InterPro | 0.772055 | KNN | F1 |
| ASR-Max | ASR-Dist | 0.892334 | KNN | Balanced_Accuracy |
| ASR-Dist | ASR-Max | 0.892334 | KNN | Balanced_Accuracy |

**Supplemental Table S3:** Statistical Analysis of Protein Representation Performance – Lysozyme C.

Dunn Test for Statistical Significance after Kruskal-Wallis Test for Different Metrics.

| Group 1 | Group 2 | P-Value | Classifier | Metric |
|---|---|---|---|---|
| InterPro | ASR-Max | 0.000363 | KNN | precision |
| ASR-Max | InterPro | 0.000363 | KNN | precision |
| ASR-Max | ASR-Dist | 0.008945 | KNN | roc_auc |
| ASR-Dist | ASR-Max | 0.008945 | KNN | roc_auc |
| InterPro | ASR-Dist | 0.011296 | KNN | roc_auc |
| ASR-Dist | InterPro | 0.011296 | KNN | roc_auc |
| InterPro | ASR-Dist | 0.067024 | KNN | precision |
| ASR-Dist | InterPro | 0.067024 | KNN | precision |
| ASR-Max | ASR-Dist | 0.356641 | KNN | precision |
| ASR-Dist | ASR-Max | 0.356641 | KNN | precision |
| ASR-Max | ASR-Max | 1 | KNN | precision |

**Supplemental Note 1**. Description and Summary of Generative VAE Architecture

### A. Summary of VAE Architecture

```
ProteinVAE(
  (encoder): Encoder(
    (conv1): Conv1d(21, 64, kernel_size=(3,), stride=(1,),\ padding=(1,))
    (bn1): BatchNorm1d(64, eps=1e-05, momentum=0.1,\ affine=True,
track_running_stats=True)
    (flatten): Flatten(start_dim=1, end_dim=-1)
    (fc1): Linear(in_features=25856, out_features=10000, bias=True)
    (fc2): Linear(in_features=10000, out_features=5000, bias=True)
    (fc3): Linear(in_features=5000, out_features=2000, bias=True)
    (fc4): Linear(in_features=2000, out_features=500, bias=True)
    (fc5): Linear(in_features=500, out_features=100, bias=True)
    (z_mean): Linear(in_features=100, out_features=100, bias=True)
    (z_log_var): Linear(in_features=100, out_features=100, bias=True)
    (sampling): Sampling()
  )
  (decoder): Decoder(
    (fc1): Linear(in_features=100, out_features=500, bias=True)
    (fc2): Linear(in_features=500, out_features=2000, bias=True)
    (fc3): Linear(in_features=2000, out_features=5000, bias=True)
    (fc4): Linear(in_features=5000, out_features=10000, bias=True)
    (fc5): Linear(in_features=10000, out_features=25856, bias=True)
    (deconv1): ConvTranspose1d(64, 21, kernel_size=(3,),\ stride=(1,), padding=(1,))
    (bn1): BatchNorm1d(21, eps=1e-05, momentum=0.1,\ affine=True,
track_running_stats=True)
    (dropout): Dropout(p=0.2, inplace=False)
  )
)
```

### B. Description of Generative VAE Architecture

GitHub: https://github.com/WoldringLabMSU/Evo-Seq

#### i. Quick mental model of "what the VAE learns" here

- The one-hot MSA-like inputs make the VAE learn family-specific patterns (including gap patterns if present).
- The KL term encourages a smooth, near-Gaussian latent space, which tends to make interpolation and sampling meaningful. Decoding produces a position × amino-acid score/probability matrix, which you then discretize into a final sequence (e.g. by argmax at each position or by sampling).

#### ii. How amino-acid sequences are encoded - "one-hot encoding" plus a 1D CNN feature extractor

**One-hot with a 21-character alphabet (20 AAs + gap)**

In the training script, each sequence is converted into a one-hot matrix of shape:

- (L, 21) where L = sequence length
- Alphabet is: `"ACDEFGHIKLMNPQRSTVWY-"` (note the final `-` gap character)

Thus, each residue position becomes a 21-dim vector with a single 1.

**Fixed-length / aligned sequences**

Note to future users… The model architecture hard-codes the flattened Conv output size as `64 * 404`, which implies the training data are expected to be length 404 (typically from an MSA with gaps).
If you feed variable-length sequences, batching will break (PyTorch cannot stack variable-length tensors without padding). For use with proteins of other lengths or variable lengths, this must be modified.

#### iii. Encoder + latent space setup - 1D CNN features + batch norm + latent dim 100

**Encoder architecture (what gets learned from sequences)**

The VAE encoder does:

1. Input batch `x`: (batch, 404, 21)
2. Transpose to channels-first: Conv: `Conv1d(21 → 64, kernel=3, padding=1)` + BatchNorm + ReLU
3. Flatten: (batch, 64*404)
4. MLP stack: `→ 10000 → 5000 → 2000 → 500 → latent_dim` (ReLU between layers)

**Latent variables (VAE core)**

The encoder outputs two vectors (size latent_dim=100):

- `z_mean` (μ)
- `z_log_var` (log $\sigma^2$)

Then it samples using reparameterization:

$$z \sim \mu + \exp(0.5 \cdot log(\sigma^2)) \odot \epsilon, \; \epsilon \sim N(0, I)$$

That is implemented explicitly in the `Sampling` module.

**Prior / regularization**

The loss includes the standard KL divergence to a unit Gaussian prior (i.e. it encourages the approximate posterior to stay near $N(0, I)$).
Thus, the "latent space setup" is: 100-D continuous Gaussian latent, regularized toward standard normal.

#### iv. How sequences are decoded (latent → amino acids)

**Decoder architecture**

The decoder mirrors the encoder in reverse:

1. Start from `z` (batch, 100)
2. MLP: `100 → 500 → 2000 → 5000 → 10000 → (64*404)`
3. Reshape to (batch, 64, 404)
4. `ConvTranspose1d(64 → 21, kernel=3, padding=1)`
5. Apply sigmoid, then transpose back to (batch, 404, 21)

**Important nuance: sigmoid + BCE (not softmax)**

The reconstruction loss is binary cross-entropy between the output tensor and the one-hot input:
That means the model is treating the output as 21 independent Bernoulli probabilities per position, rather than a categorical distribution with softmax.

Note to future EvoSeq users… In practice, to turn the decoder output into an amino-acid sequence, you typically:

- take an argmax over the 21 channels at each position, or
- renormalize per position and sample.

Thus, users could consider a simple decoder-to-string function (based on the amino acid + gap alphabet) that would look like:

```
import torch
AA = "ACDEFGHIKLMNPQRSTVWY-"
def decode_logits_to_seq(y):
    # y: (L, 21) in [0,1] from sigmoid
    idx = torch.argmax(y, dim=-1).tolist()
    return "".join(AA[i] for i in idx)
```

Then you would usually post-process gaps (-) depending on whether you want an *aligned* sequence (keep them) or an *ungapped* protein sequence (strip them or remove columns).

#### v. How generation actually happens in this project

EvoSeq captures two common workflows:

1. **Unconditional sampling from the prior**
   Sample $z \sim N(0, I)$, decode, argmax → sequence.
2. **Sampling around training examples ("generate from training")**
   Encode a real sequence to get (μ, log($σ^2$)), sample $z \sim N(\mu, \sigma^2)$, decode.
   (Our EvoSeq README points to a "generate from training" script path, even though that exact filename may differ across projects.)

Training details in the script:

- shuffles FASTA sequences, one-hot encodes, 90/10 train/val split
- Adam lr=1e-4, weight_decay=1e-8, up to 300 epochs, early stopping patience=5, batch_size=16
