## Supplementary Methods for "EvoSeq-ML: Advancing Data-Centric Machine Learning with Evolutionary-Informed Protein Sequence Representation and Generation"

### BAli-Phy replicate workflow, ASR generation, and AP-LASR integration

This document provides step-by-step instructions to reproduce the BAli-Phy replicate workflow used in this study, generate the MAP tree and consensus alignment, produce an ASR.state file for downstream analyses, and run the modified AP-LASR script. An additional cleanup/finalization script is included to repair common tree/alignment naming and ordering issues prior to downstream use (EvoSeq).

### SI 1. Files Provided

**AP-LASR.py**: Modified AP-LASR Python script used for ASR and downstream AP-LASR steps.

**bp_aplasr_bali-phy4.1_full_pipeline.sb**: Primary end-to-end SLURM pipeline (FASTA → 5 BAli-Phy runs → select 3 → bp-analyze → MAP.tree/consensus alignment → ASR.state → AP-LASR).

**evoSeq_cleanup_finalize_ASRstate.sb**: Optional cleanup/finalization script to rebuild/sanitize `MAP.tree` and consensus alignment and regenerate `ASR.state` from existing BAli-Phy runs.

**Species_Name_Truncator_For_PAML.py**: Optional Windows helper to truncate FASTA headers to 10 characters for strict PHYLIP/PAML workflows and generate an Excel mapping (original → truncated).

### SI 2. Requirements and Environment

Compute environment (recommended):

- HPC cluster with a SLURM scheduler.
- Singularity installed and available on compute nodes.
- BAli-Phy Singularity image available in `$HOME` (see below).

Modules used in the provided SLURM scripts (may vary by cluster): `GCC/12.3.0` and `OpenMPI/4.1.5`. The main pipeline also loads `Biopython`, `MAFFT`, `CD-HIT`, `IQ-TREE`, and `matplotlib` for downstream AP-LASR tasks. Adjust module names/versions as needed for your environment.

### SI 3. Bali-Phy Singularity Image

The workflow runs BAli-Phy tools via Singularity. Place the following image in `$HOME` (`/mnt/home/<user>/`):

- `bali-phy-4.1-linux64-intel-singularity.sif`

If the image filename or location differs, update the SIF path (or the singularity exec lines) in the SLURM scripts.

### SI 4. Quick Start

1. Copy your input protein FASTA into your working directory on the cluster (same directory as the `.sb` scripts).
2. Confirm the BAli-Phy Singularity image is present in `$HOME`.

3. Submit the main pipeline: `sbatch bp_aplasr_bali-phy4.1_full_pipeline.sb`
4. Key outputs will be written to `bp_analyze_output/` (including `MAP.tree`, consensus alignments, and `ASR.state`).

### SI 5. Main pipeline details (bp_aplasr_bali-phy4.1_full_pipeline.sb)

Overview of steps executed by the main pipeline:

1. Detects an input FASTA file in the working directory.
2. Optionally cleans FASTA headers into `cleaned_input.fasta` by keeping only the first token of each header (recommended to avoid downstream name parsing issues).
3. Launches five independent BAli-Phy MCMC replicates from the same input FASTA.
4. After burn-in, extracts posterior values from each replicate and computes the median posterior for each run.
5. Selects the three runs with the most similar post burn-in median posterior values to reduce sensitivity to outlier chains prior to downstream summarization.
6. Runs `bp-analyze` on the selected replicates to produce a consensus summary and the maximum a posteriori (MAP) tree (`MAP.tree`) in `bp_analyze_output/`.
7. Reorders the consensus alignment to match the `MAP.tree` tip order (`alignment-cat`).
8. Runs `summarize-ancestors` to generate bp_analyze_output/ASR.state for downstream AP-LASR analyses.
9. Generates `posterior_medians.png` (boxplot of posteriors across replicates).
10. Runs AP-LASR using `MAP.tree` and the input FASTA to generate downstream outputs.

### SI 6. Expected inputs and outputs

Inputs:

- Input protein FASTA file (`*.fasta`) in the working directory.
- BAli-Phy Singularity image (`.sif`) in $HOME.
- `AP-LASR.py` in a known path (the provided pipeline calls /`mnt/home/k0099424/AP-LASR.py`; update this path for other users).

Primary outputs (generated by the pipeline):

- `bp_analyze_output/MAP.tree` - MAP tree produced by bp-analyze.
- `bp_analyze_output/P1.consensus.pd-multiply.fasta` - consensus alignment reordered to match `MAP.tree`.
- `bp_analyze_output/ASR.state` - ancestral sequence reconstruction state file generated by summarize-ancestors.

- `bp_analyze_output/bp_analyze.log` - bp-analyze log output.
- `posterior_medians.png` - boxplot of posterior distributions across the five BAli-Phy runs.
- BAli-Phy run directories (e.g., `cleaned_input-1` ... `cleaned_input-5`) containing `C1.log`, `C1.P1.fastas`, and `C1.trees`.

#### SI 7. Cleanup/finalization (evoSeq_cleanup_finalize_ASRstate.sb)

This script is intended for cleanup/finalization and troubleshooting, not for running EvoSeq directly. It is useful if MAP.tree/consensus alignment ordering or taxa naming issues prevent downstream analyses.

What it does:

1. Finds at least three BAli-Phy run folders (prefers `-1`, `-3`, `-5` if present).
2. Uses an existing `bp_analyze_output/MAP.tree` if present; otherwise builds a consensus tree from the selected runs' `C1.trees`.
3. Builds an unordered consensus alignment from the selected runs (cut-range → alignment-chop-internal → alignment-max).
4. Sanitizes tip names in `MAP.tree` by removing trailing underscores at delimiters and writes `bp_analyze_output/MAP.tree.sanitized`.
5. Reorders the consensus alignment to match the sanitized tree (`alignment-cat`).
6. Performs a taxa-set check between the tree tips and the alignment headers and prints differences.
7. Regenerates `bp_analyze_output/ASR.state` using summarize-ancestors on the ordered alignment and sanitized tree.

#### SI 8. Optional: PAML/PHYLIP taxa-name truncation **(Species_Name_Truncator_For_PAML.py)**

Some strict PHYLIP/PAML workflows require short (≤10 character) taxa names. This optional script runs on Windows to:

- Create a FASTA file with truncated headers (first 10 characters) suitable for strict PHYLIP conversion.
- Generate an Excel mapping of original → truncated names for traceability.

Note: Verify that truncation does not create duplicate IDs; duplicates must be resolved before running phylogenetic tools.

### SI 9. Troubleshooting

BAli-Phy run folders are not detected

- Ensure run directories follow the expected pattern (`cleaned_input-1 … cleaned_input-5 or Final_Sequences_Short-1 …`), or place run directories in `$HOME` with `C1.P1.fastas` and `C1.trees` present.

Singularity image not found

- Place the `.sif` image in `$HOME` or update the SIF path in the SLURM scripts.

AP-LASR.py path differs

- Update the python command in `bp_aplasr_bali-phy4.1_full_pipeline.sb` to point to the correct script path.
